# Somatic Transthyretin-Related Proteins in C. elegans Govern Reproductive Longevity by Sustaining Sperm Integrity and Timely Ovulation

**DOI:** 10.1101/2024.09.13.612966

**Authors:** Tingshan Wu, Haochen Lyu, Zhao Wang, Zhaoyang Jiang, Yingchuan B. Qi

## Abstract

The decline in reproductive capability during adult life is critical for health, but its mechanism is elusive. We systematically analyzed the developmental role of an expanded TTR family of proteins, structurally analogous to mammalian thyroid hormone-transporting Transthyretin, and identified three paralogous proteins, TTR-15, TTR-16, and TTR-17, differentially expressed in somatic cells of the gonads and secreted around gametes in *C. elegans*. Simultaneous inactivation of TTR-15, TTR-16, and TTR-17 leads to a rapid reduction in reproductive capacity in middle age. While oocyte and sperm production remain unaffected in the triple mutants, late-onset infertility results from stalled ovulation. Mechanistically, the absence of TTR-15, TTR-16, and TTR-17 causes sperm to prematurely deplete the cytoplasmic pool of major sperm protein (MSP), released via non-conventional vesicle budding as a signal for ovulation. We propose that the somatic gonads play a central role in maintaining sperm integrity post-production and determining the duration of the reproductive age.

**Highlights:**

- Systematic analysis of TTR family proteins reveals diverse expression and critical functions.
- TTR-15, TTR-16, and TTR-17 are secreted around gametes in *C. elegans*.
- TTR-15/16/17 triple KO exhibits middle-age onset infertility due to stalled ovulation.
- MSP, a signal for ovulation, is prematurely depleted from sperm in the absence of TTR-15/16/17.

## Introduction

Developmental timing control is fundamental to an organism’s fitness, guiding events spanning embryogenesis, juvenile growth, maturation, and aging^1–3^. One crucial aspect of developmental timing involves the reproductive decline^4,5^, which is conceivably tightly linked to the overall life cycle schedule. *C. elegans* has been a pivotal model for studying developmental timing and aging^6,7^. In *C. elegans*, both hermaphrodites and males exhibit a reproductive peak lasting a few days upon reaching adulthood, followed by a rapid decline, after which they continue to live for an extended adult lifespan^8,9^. Although classical genetic analysis has revealed miRNA-triggered pathways that guide stage transitions in larvae, the mechanisms that pace adult development remain poorly defined.

*C. elegans* hermaphrodites produce both sperm and ova for self-fertilization, while males produce only sperm and the crossing with hermaphrodites introduces genetic diversity^10,11^. During larval development, hermaphrodites produce sperm (∼150 each gonad arm) initially and switch to continuous oogenesis in adulthood^12^. Mature oocytes are fertilized as they pass through the spermatheca, where sperm are stored (**Figure S1A**).

Gametes of *C. elegans* undergo complex maturation similar to mammals^13–15^. Primary oocytes arrest at prophase of the first meiotic division and are associated with somatic gonad sheath cells. Major sperm protein (MSP) released by sperm triggers oocyte maturation and ovulation^16^. Without sperm, ovulation stalls, and arrested oocytes accumulate in the oviduct^17^.

Spermatogenesis produces round spermatids with condensed nuclei, mitochondria, and membranous organelles (MO)^18^. In hermaphrodites, spermatids activate when pushed into the spermatheca by ovulating oocytes. In males, spermatid activation occurs upon ejaculation. Activated spermatids form motile spermatozoa via MSP polymerization^19,20^; therefore, MSP plays a dual role in sperm motility and oocyte maturation, linking these processes for reproductive success.

MSP is a cytoplasmic protein unrelated to actin and lacks a secretory signal sequence. The polymerization of MSP drives the formation of pseudopods, enabling spermatozoa to crawl^19,20^. As an extracellular signal, MSP is released by sperm through the unconventional budding of double-membraned vesicles, though the mechanism is unclear^21^. MSP is released by both immotile spermatids (in oviducts) and motile spermatozoa (in the spermatheca), with differences suggesting controlled MSP vesicle release, despite the underlying mechanism remaining elusive.

Interactions between germ and somatic components in the reproductive system are complex and essential for proper function^22^. In *C. elegans*, gonad sheath cells support the germ cell proliferation, gametes differentiation, and meiotic maturation^23^. They also instruct oocytes to release F-series prostaglandins (PGs) to guide sperm chemotaxis^24^. The spermatheca provides the necessary environment for sperm activation, as immotile spermatids activate upon entry, though the mechanism is unclear^25–27^. This sperm-somatic interaction in *C. elegans* may parallel mammalian sperm maturation and capacitation, regulated by factors from the epididymis, seminal vesicles, prostate gland, and female reproductive ducts, which are also not well understood^28,29^.

While studying the role of the transmembrane transcription factor MYRF during larval stages^30^, we consistently noticed that a family of genes encoding transthyretin-related proteins (TTR) are differentially expressed in development-aberrant mutants. Motivated by transthyretin’s role in binding thyroid hormones and retinol^31^—key regulators of development—we systematically analyzed TTR expression and function in *C. elegans*. We found that three close paralogs, TTR-15, TTR-16, and TTR-17, are differentially expressed in the somatic gonad and secreted around gametes. Deletion of these genes significantly decreases progeny; most notably, the triple mutants exhibit a middle age-onset ovulation block. Our analysis indicates that the absence of these proteins accelerates sperm quality decline, marked by premature depletion of sperm MSP. To our knowledge, this is the first report of genetic mutants specifically affecting MSP release.

Additionally, inactivation of TTR-15, TTR-16, and TTR-17 results in varied developmental timing, with some mutants showing delayed development and others accelerated growth from L4 to adult. Together with defects in middle-aged sperm, these findings suggest that TTR genes act as novel regulators of developmental timing and reproductive function.

## Results

### Diverse Tissue Expression Patterns of TTR Family Genes in C. elegans

Transthyretin was first recognized in the 1940s for its function in binding thyroid hormones and retinol. With advancements in genomics, transthyretins were later discovered to belong to a larger family known as transthyretin-like proteins (TLPs) or transthyretin-related proteins (TRPs). These proteins are found in a wide range of organisms, from bacteria to plants, indicating their extensive evolutionary presence. The genome of C. elegans contains an expanded family of proteins (59 members) with sequence homology to the transthyretin, which has been designated as transthyretin-related proteins (TTRs) (**Figure 1, 2A**). The expression and function of the TTR family genes are largely unexplored.

**Figure 1.**
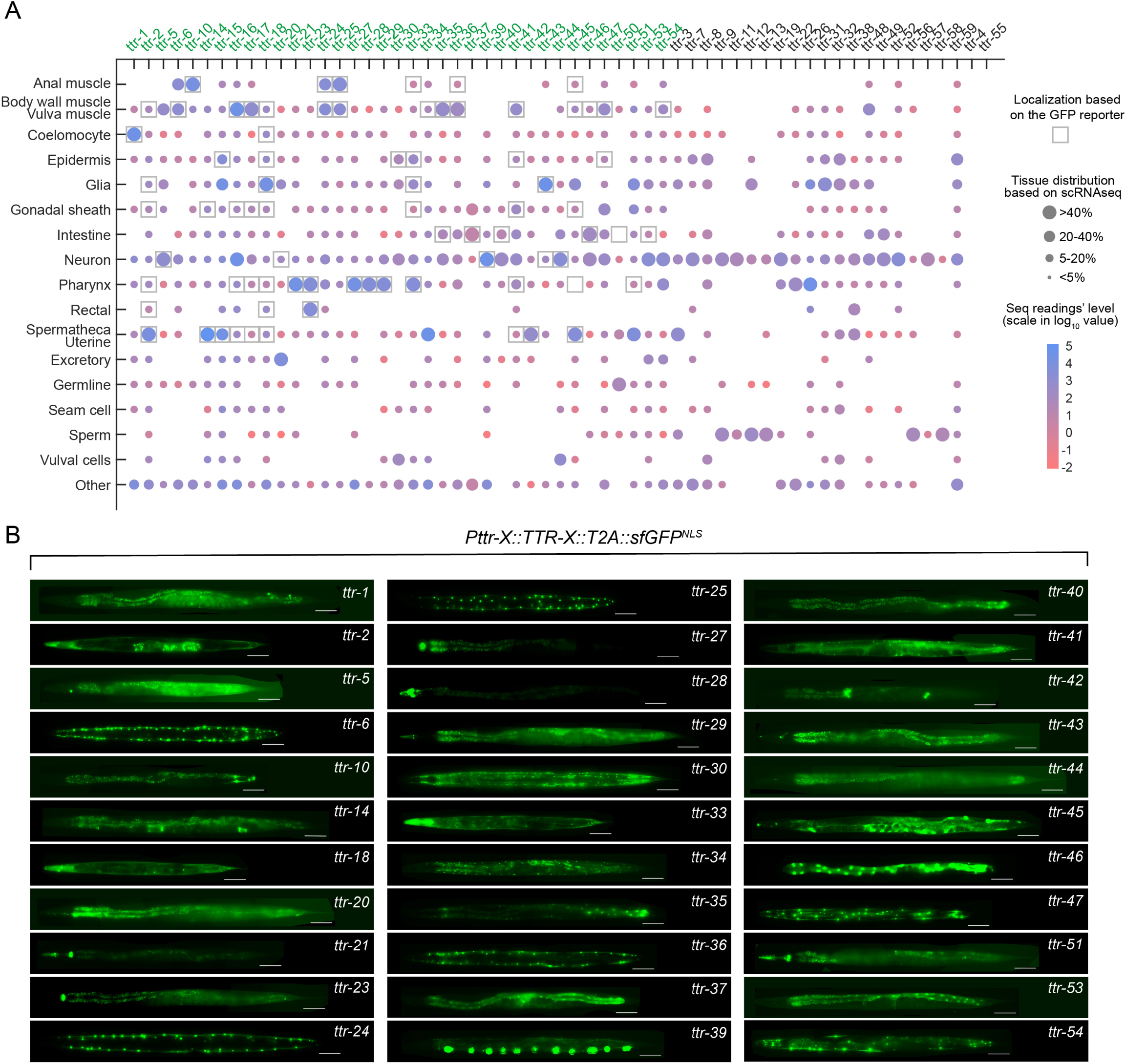
Systematic Analysis of Expression of TTR Family Genes in C. elegans. A) This chart maps the tissue distribution of each TTR gene. Round dots reflect tissue enrichment from single-cell RNA sequencing analysis (data from the CeNGEN consortium). Dot size represents the percentage of transcript counts in specific tissues, while color coding indicates standardized expression levels. Boxes signify the detection of TTR through transcript analysis in transgenic lines, with gene names highlighted in green. B) Representative images showcase the expression of TTR genes using either extrachromosomal or integrated tandem array transgenes. Each transgene construct includes the TTR gene’s promoter, exons, and introns, and also has a T2A::GFP::NLS sequence fused to the C-terminus of the TTR ORF.

Our phylogenetic analysis of the 59 TTR family genes in C. elegans reveals their conservation and diversification, indicating the existence of multiple TTR subfamilies (**Figure 2A**). The majority of these genes contain secretory peptide signals, although a few members (ttr-40, ttr-56 and ttr-57) may lack conspicuous signals.

**Figure 2.**
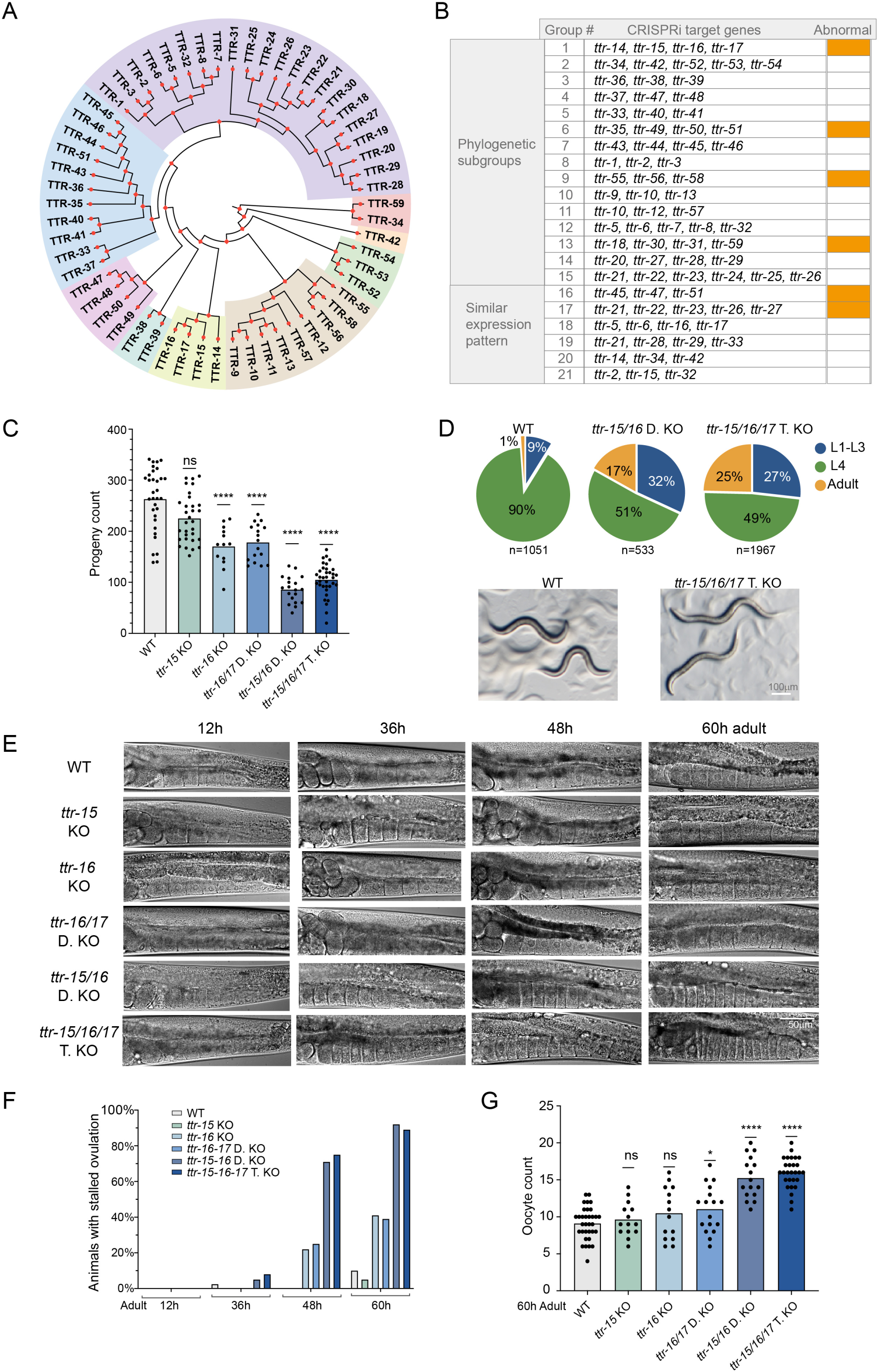
TTR-15, TTR-16, and TTR-17 Triple Knockout Animals Show Late-Onset Ovulation Defects. A) A phylogenetic tree indicating the existence of multiple TTR subfamilies. B) This summary presents the results of multi-targeted CRISPRi groups based on their phylogenetic subgroups or similarities in tissue expression patterns. We visually examined the general phenotype of transgenic F1 generations, focusing primarily on body morphology and viability. Groups labeled as ‘abnormal’ displayed a phenotype penetrance exceeding 50% among the transgenic F1 individuals. C) Significant reduction in progeny number in ttr-15/16/17 triple KO animals. The graph quantifies the total progeny counts for ttr-15, ttr-16, ttr-17, and compound mutants. Refer to supplemental S1 for progeny counts from tracking individual animals. D) Abnormal developmental timing in ttr-15/16 double KO and ttr-15/16/17 triple KO animals. Larvae were synchronized and observed until 40 hours, by which time most wild-type animals reached the L4 stage. In contrast, triple KO animals exhibited varied developmental timing—some delayed, others precocious. The majority of the delayed group eventually matured into normal young adults. On the right are representative images of wild-type L4 and more advanced triple KO young adults. E) ttr-15/16/17 triple KO animals show late-onset ovulation defects. In wild-type animals, oocyte maturation and fertilization continue up to 3-4 days. Triple KO animals show a normal pattern of oocyte arrangement in the oviduct during the first day of adulthood. However, by the second day, ovulation decreased while oocyte production continued, leading to an accumulation and compression of oocytes in the oviduct. F) The proportion of animals that show stacked oocytes in the oviduct over the reproductive period. The proportion of ovulation-defective animals significantly increased during the second day in triple KO animals. G) Counting oocytes in one gonad arm in the third day adults.

To explore the tissue-specific expression of these TTR genes, we generated expression reporter transgenes for 30 TTR genes (**Figure 1**). Each transgene incorporated the respective TTR gene’s promoter, exons, and introns, and the 3’ untranslated region from unc-54. A T2A peptide sequence and an NLS-carrying GFP are placed before the stop codon. This setup allows GFP to be co-expressed with TTR via the T2A peptide, mirroring endogenous TTR expression.

Our studies have shown that TTR genes are expressed in a variety of specific tissues, including the epidermis, certain neurons, muscles, the intestine, reproductive organs, etc. While some TTR genes are broadly expressed across these tissues, others show a more limited distribution, suggesting diverse physiological roles. We created a matrix chart for tissue expression specificity by comparing previously reported single-cell RNA sequencing (scRNAseq) data with our transgene reporter analyses (**Figure 1A**). We found that the reporter results generally confirm the expression sites with highly enriched transcripts identified by scRNAseq. However, the reporter signals exhibit a much more restricted pattern than the scRNAseq data suggests. For example, ttr-39 has been observed in several studies to be highly specific to DD and VD neurons (also shown in **Figure 1B**), but the scRNAseq data suggests a much broader expression. Therefore, our reporter analysis provides useful information to further delineate the function of TTR family genes.

### Systematic CRISPRi Analysis Reveals Critical Functions of TTR Genes

Considering the impact of potential gene redundancy, we employed CRISPR interference to target several TTR genes simultaneously to explore their roles in C. elegans (**Figure 2B**). For each targeted TTR gene, two guide RNAs (gRNAs) were designed to target its promoter region. Given the exploratory nature of the analysis and the partial penetrance of the technique, we did not follow strict selection criteria, but generally group the candidates based on their subfamily affiliations or expression pattern similarities. The gRNA expression vectors were pooled and used for transgenic microinjection to disrupt 3-5 TTR genes concurrently.

The resulting phenotypes in the F1 generation from these gene disruptions varied, ranging from superficially wild-type to developmental lethality, morphological abnormalities, and reproductive deficiencies. These findings suggest critical functions for TTR genes in developmental and reproductive processes in C. elegans, though their specific roles remain to be further defined.

### Reduced Fecundity and Altered Developmental Tempo in TTR Triple Mutants

We decided to further investigate the function of TTR-15, TTR-16, and TTR-17, as these genes were among those for which CRISPRi caused consistent abnormalities. We generated gene deletion mutants for the three TTR using CRISPR-Cas9 method (**Figure S1C**). TTR-16 and TTR-17, being genetically linked, were deleted both individually and simultaneously, while TTR-15, located on a separate chromosome, was deleted individually. While single-gene mutants showed no noticeable phenotype, double mutants of TTR-15 and TTR-16, as well as triple knockouts of TTR-15, TTR-16, and TTR-17, exhibited a significant decrease in progeny (**Figure 2C**; **Figure S1D**).

To assess the general development of these mutants, we tracked the stage progression of TTR triple KO animals from synchronized L1 larvae. Notably, TTR triple KO mutants exhibited marked asynchronous developmental paces (**Figure 2D**). By 40 hours at 20°C, a time when most wild-type animals reach the L4 stage, some of the triple KO population was delayed at the L2-3 stages, whereas a considerable portion had accelerated development, reaching adulthood earlier. While developmental delays due to deficiencies in biological processes are not uncommon, accelerated development is unusual and a few instances are associated with altered nutrient gradients in the *C. elegans* diet. This abnormal stage variability observed in triple KO mutants suggests the significant impact of TTR-15, TTR-16, and TTR-17 in regulating growth and developmental pacing.

### Late-Onset Oocyte Maturation Block in TTR Triple mutant

In addition to reduced progeny in triple KO mutants, a notable abnormality in these animals is the presence of stacked oocytes in the proximal gonadal arm (**Figure 2E**), a phenotype similar to that observed in *fog-2* mutants, which lack functional sperm. Normally, sperm release MSP to initiate oocyte maturation and ovulation, involving cytoskeletal remodeling, detachment of oocytes from sheath cells, and a shape change from cylindrical to oval^14,32–34^. Without normal ovulation, oocytes remain trapped in the oviduct.

Distinct from any other known sperm-defective mutants that show stacked oocytes in the oviduct from early adulthood, the oocyte maturation block in triple KO mutants is late-onset (**Figure 2E, F**; **Figure S1B, D**). Ovulation appears normal during the first 24 hours of adulthood but starts to show defects from the second day onward, with severe oocyte stacking by the third day of adulthood. This maturation block is also manifested by the cortical enrichment of microtubules in triple mutant oocytes, in contrast to the even distribution throughout the cytoplasm observed in control animals (**Figure S1E**). Comparing the severity of the oocyte defects among single, double, and triple mutants, it appears that TTR-15 and TTR-16 together play a predominant role, while TTR-17 plays a minor role (**Figure 2C-G**). This late-onset ovulation deficiency suggests that the absence of TTR-15, TTR-16, and TTR-17 disrupts a specific adult reproductive schedule, affecting the normal continuation of oocyte maturation and ovulation.

### Sperm Persist but Are Incompetent in Middle-Aged TTR Triple KOs

The stalled ovulation suggests depleted sperm in the triple mutants, similar to how ovulation ceases in aged wild-type hermaphrodites after their sperm are used up. Surprisingly, investigating day 2-3 triple mutants revealed a significant presence of sperm in the spermatheca (**Figure 3A, B**; **Figure S2A**), challenging the assumption that a lack of sperm causes failed oocyte maturation. This suggests that the lack of ovulation could be due to defective oocytes or somatic gonad cells.

**Figure 3.**
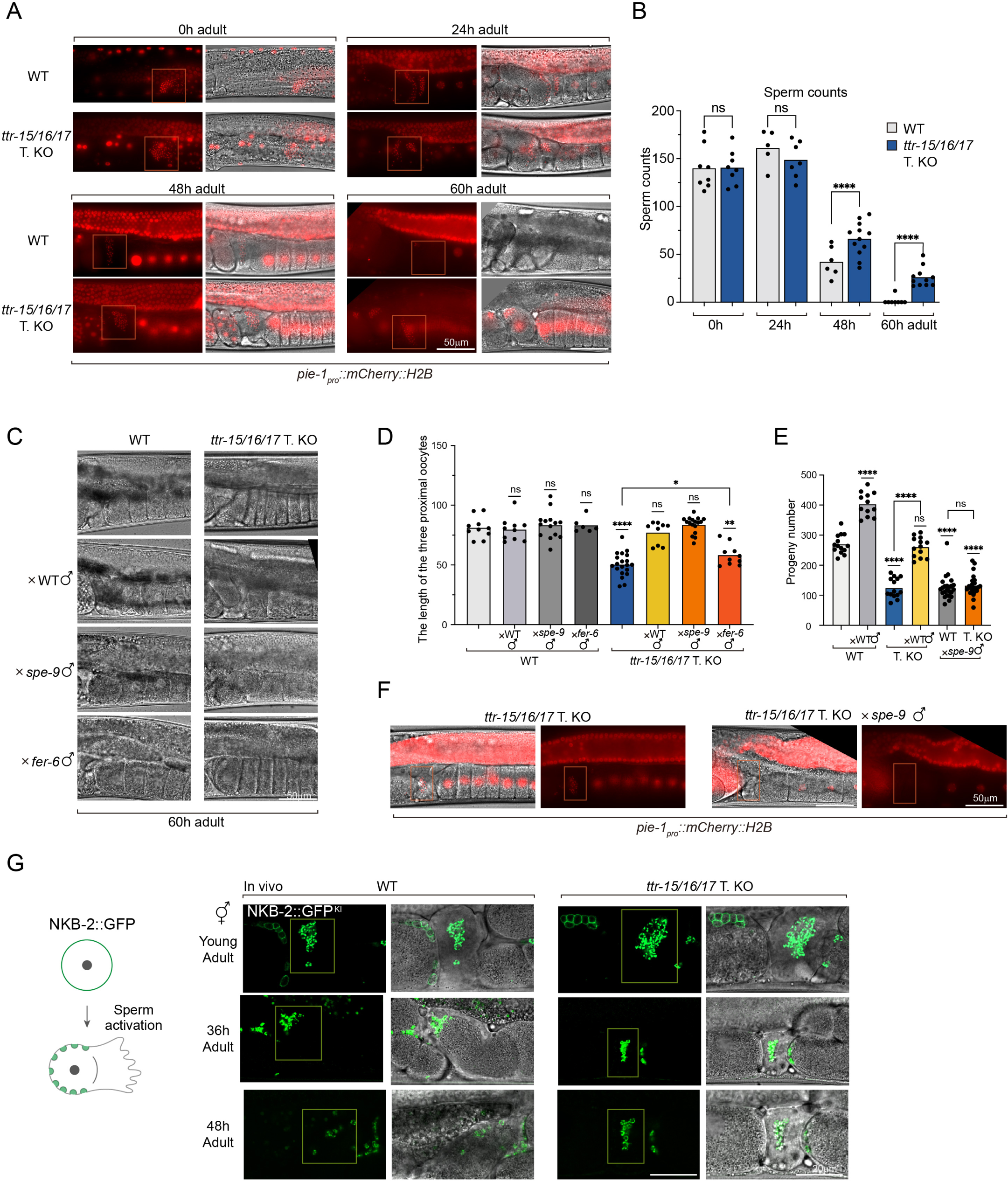
Stalled Ovulation in Middle-Aged TTR-15/16/17 Triple KO Animals Linked to Dysfunctional Sperm. A) Many sperm are present in triple KO animals by day 2-3 when ovulation stalls. These sperm persist in the spermatheca of triple KO animals. In contrast, wild-type sperm are used up by day 3-4 through fertilization. The nuclei of sperm and oocytes are labeled by mCherry::H2B expressed in the germline. The red rectangle labels the position of the spermatheca in which small, dense nuclei of sperm can be detected. B) Sperm counts in wild-type and triple KO animals over the reproductive period. C) Ovulation defects in middle-aged triple KO animals are corrected by crossing with wild-type males or spe-9 males. spe-9 sperm are motile but fertilization-defective. The ovulation defects in triple KO are partially corrected by crossing with fer-6 males. fer-6 sperm are defective in sperm activation and are immotile. D) Quantification of the length of the three most proximal oocytes, as described in C. The decrease in length corresponds with the stacked oocyte or the ovulation block. E) Progeny counts when triple KO animals are crossed with wild-type males or spe-9 mutant males. The progeny number is significantly increased by crossing with wild-type males. The presence of spe-9 male sperm antagonizes the wild-type sperm, thereby decreasing the progeny number in wild-type animals. spe-9 sperm did not further reduce nor increase the progeny number in triple KO animals. F) Representative images of triple KO animals after they are crossed with spe-9 males. The nuclei are labeled by mCherry::H2B. Ovulation is restored in triple KO animals. The self-sperm of triple KO animals disappeared, presumably pushed out by the passing oocytes. The disappearance of self-sperm in triple KO animals suggests they have defective motility. G) Sperm activation is normal in ttr-15/16/17 triple KO animals. On the left, the drawing illustrates the change in distribution patterns of NKB-2::GFP during the transition from spermatids to spermatozoa. Sperm activation appears normal in triple KO based on NKB-2::GFP patterns. Activated spermatozoa in the spermatheca are highlighted in boxes. Spermatids in the oviduct have GFP signals only on the cell membrane.

To test this notion, we supplied the triple mutant hermaphrodites with exogenous sperm by crossing them with wild-type males (**Figure 3C-E**). This cross significantly increased progeny numbers and restored ovulation, indicating that the oocytes and gonad sheath cells in triple KOs are relatively healthy.

The defective ovulation despite the presence of sperm in triple KOs suggests that these sperm may be abnormal, particularly in releasing MSP signals. We crossed triple KO hermaphrodites with spe-9 mutant males, whose sperm are motile but fertilization-defective^35^, and the exogenous spe-9 male sperm successfully rescued ovulation (**Figure 3C-E**). The ovulation defect in triple KOs was partially rescued by crossing with fer-6 mutant males, whose sperm cannot be normally activated and are therefore mostly immotile^26^. This evidence pinpointed the issue to sperm-derived signaling. Notably, despite restored ovulation, the oocytes failed to be fertilized by either the spe-9 male sperm (which is expected) or the self-sperm in triple KO hermaphrodites, as the progeny numbers were not improved (**Figure 3E**). This suggests overall poor competence of these self-sperm from middle age onwards. In fact, those stored self-sperm were rapidly depleted after the cross (**Figure 3F**), implying their deficient motility in these middle-aged sperm.

### Normal Spermatid Activation in TTR Triple Mutants

Spermatids are spherical and non-motile; they transform into mature, motile spermatozoa through “sperm activation” or “spermiogenesis,” which involves membrane organelle (MO) fusion cortically to remodel the cell surface and the formation of pseudopods for movement (**Figure S1A**). To determine if sperm activation occurs normally in triple mutants, we analyzed the distribution of NKB-2, a sodium-potassium pump protein that shifts from being evenly distributed across the spermatid membrane to being concentrated at MO fusion sites upon activation^36^. In TTR-15, TTR-16, and TTR-17 triple KO hermaphrodite sperm, NKB-2::GFP showed normal activation patterns, characterized by circular signals on unactivated spermatids in the oviduct and discrete-patched pattern in activated sperm in the spermatheca (**Figure 3G**).

We also examined another marker protein, SPE-38::mCherry^37^, which relocates from intracellular vesicles to the cell membrane upon activation, in triple KO mutants. Due to the weak intensity of SPE-38::mCherry signals in live animals, we examined in-vitro activated sperm from young triple knockout males (**Figure S2B, C**). These sperms were similar in morphology and SPE-38::mCherry distribution to wild-type male sperm. The sperm of triple KO male animals were prematurely activated by the *swm-1* mutation, similar to control animals, suggesting that the sperm of triple KO mutants can be activated in vivo (**Figure S2D**). Together, the initial normal fertility and correct distribution pattern of NKB-2 and SPE-38 suggest that the transformation to motile spermatozoa occurs normally in young-aged triple mutants.

### MSP Filaments may Assemble When Motile and Disassemble When Stationary

As MSP is the key signal to induce ovulation, we investigated the levels of MSP in the persistent sperm of middle-aged triple mutants. MSP is abundant in *C. elegans* sperm, and there are 47 MSP paralogs in the genome^38^. We generated a GFP tag knock-in line for one of the MSP paralogs but did not observe discernible signals in the animals. Therefore, we immunostained dissected hermaphrodite gonads using an MSP antibody.

The MSP signals exhibited a pattern consistent with previous descriptions (**Figure S3A**; **Figure 4A, C**). In spermatocytes, they formed multiple, large, shallow crescent-shaped concentrations, enclosed in fibrous bodies (FB). During the maturation of spermatids, MSP appears to be released from FBs, becoming more diffused into the cytoplasm.

**Figure 4.**
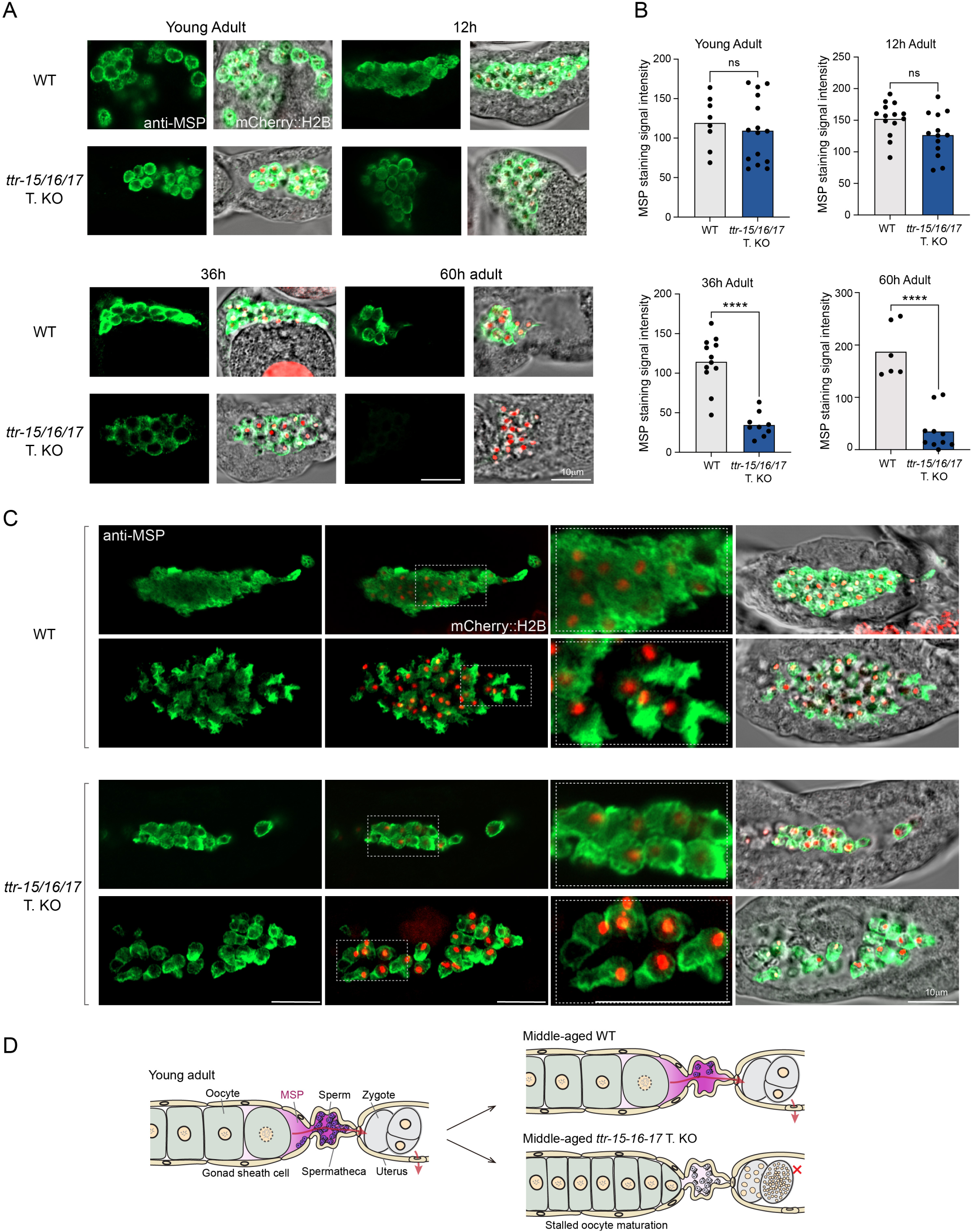
Premature Depletion of MSP in Sperm in Middle-Aged TTR-15/16/17 Triple KO Animals. A) Immunostaining of MSP on the dissected gonads of wild-type and triple KO animals. The stage-matched wild-type and mutant animals are processed in parallel. The intensity of MSP staining is comparable between wild-type and mutants in young adults. The MSP signal intensity significantly decreases in mutants on the second day and becomes undetectable in mutants on the third day. Refer to the supplemental figure S3 for the staining of genotype-matched groups. B) Quantification of the intensity of MSP signals from staining as in A. Wild-type animals usually exhaust their stored sperm by the end of third day into adulthood, making wild-type animals with retained sperm rare. C) Immunostaining of MSP reveals two types of localization patterns that may be linked to the state of sperm crawling activity and MSP polymerization. The top row of wild-type and triple mutants shows the aggregation state, in which pseudopod projections are not obvious and MSP is enriched in the cytoplasm. The bottom row of wild-type and triple mutants shows the dispersed state, in which sperms exhibit characteristic pseudopod extensions with concentrated MSP signals in pseudopods, appearing in a filament-like pattern. MSP signals are frequently absent in the pseudopods of sperm in triple mutants, resulting in an empty bag appearance. D) The illustration summarizes that stalled oocyte maturation in middle-aged ttr-15, ttr-16, and ttr-17 triple KO mutants results from the premature depletion of Major Sperm Protein (MSP) in sperm. Despite the presence of sperm, the failure to release MSP leads to impaired oocyte maturation.

Immunostaining of sperm in situ in the spermatheca reveals two distinct MSP distribution patterns in spermatozoa (**Figure 4C**). Previously reports of in vitro staining and imaging suggest that activated spermatozoa maintain a polarized morphology with extended pseudopods, where MSP is concentrated. However, this is not always observed in our in vivo preparations. In many immunostained gonads, sperm showed even MSP distribution, similar to unactuated spermatids, without polarized MSP localization or obvious pseudopod protrusion (**Figure 4C**). These sperm were often packed together in the spermatheca.

Conversely, in some gonads with more dispersed sperm, many exhibited distinct pseudopod extension with prominent MSP signals (**Figure 4C**), suggesting extensive MSP filaments. This correlation between pseudopod extension and sperm dispersion implies that MSP polymerization in pseudopods is dynamic and regulated as needed. When sperm disperse, MSP assembles into filaments for motility. When sperm pack in the spermatheca, MSP filaments disassemble, possibly to conserve energy and avoid unnecessary movement.

### Premature Depletion of MSP in TTR Triple Mutant Sperms

For analyzing MSP levels through staining, it is essential to process tissues from age-identical wild type and triple KO animals in parallel. Since it is impractical to process all age groups at once, we devised two control schemes: processing genotype pairs of the same age in parallel (**Figure 4A, B**), and processing age groups of the same genotype (**Figure S3B**). These schemes produced consistent results.

MSP levels between wild type and triple KO animals were comparable at the L4 and 12-hour adult stages. A mild decrease in MSP levels was observed in both wild type and triple KO animals at 24 hours, with a slightly lower level in the triple mutants, though not statistically significant. In 2-day-old adults, MSP signals significantly decreased in mutants and became barely detectable in 3-day-old adults. This rapid decline in MSP in mutant sperm explains the normal ovulation in young mutant animals but stalled ovulation in middle-aged animals.

Two distinctive MSP staining patterns were observed in triple KO sperm. Cytoplasmic diffused MSP correlated with packed spermatozoa, while intensive MSP concentration away from the nucleus was associated with crawling spermatozoa with extended pseudopods (**Figure 4C**). In middle-aged triple mutants, sperm pseudopods often lacked internal MSP signals, though the cortex retained sufficient MSP. This “empty-bag” pattern contrasts with control animals, where pseudopods exhibited intensive MSP signals (**Figure 4C**). This observation aligns with the premature drop in MSP levels in mutant sperm.

The results suggest that MSP for release as extracellular vesicles and for polymerization into filaments comes from the same cytoplasmic pool. As this pool shrinks, there is insufficient MSP for pseudopod polymerization or extracellular signaling. This expedited decrease in MSP levels in mutant sperm impacts both the signaling strength and crawling capacity (**Figure 4D**).

### Differential Secretion of TTR-15, TTR-16, and TTR-17 from Somatic Gonad Cells

To visualize their expression and localization in vivo, we generated GFP-tagged lines for TTR-15, TTR-16, and TTR-17 (**Figure 5A, B**; **Figure S4B**). Imaging of these GFP-tagged lines revealed differential expression patterns: TTR-15::GFP was primarily found in the lumen of the spermatheca and uterus, with concentrations around the spermatozoa, although it did not tightly bind to sperm. TTR-16::GFP exhibited much more intense signals than TTR-15::GFP and was localized broadly in the pseudocoelom, surrounding the pharynx, and in the lumen of the oviduct, spermatheca, and uterus. TTR-17::GFP was undetectable.

**Figure 5.**
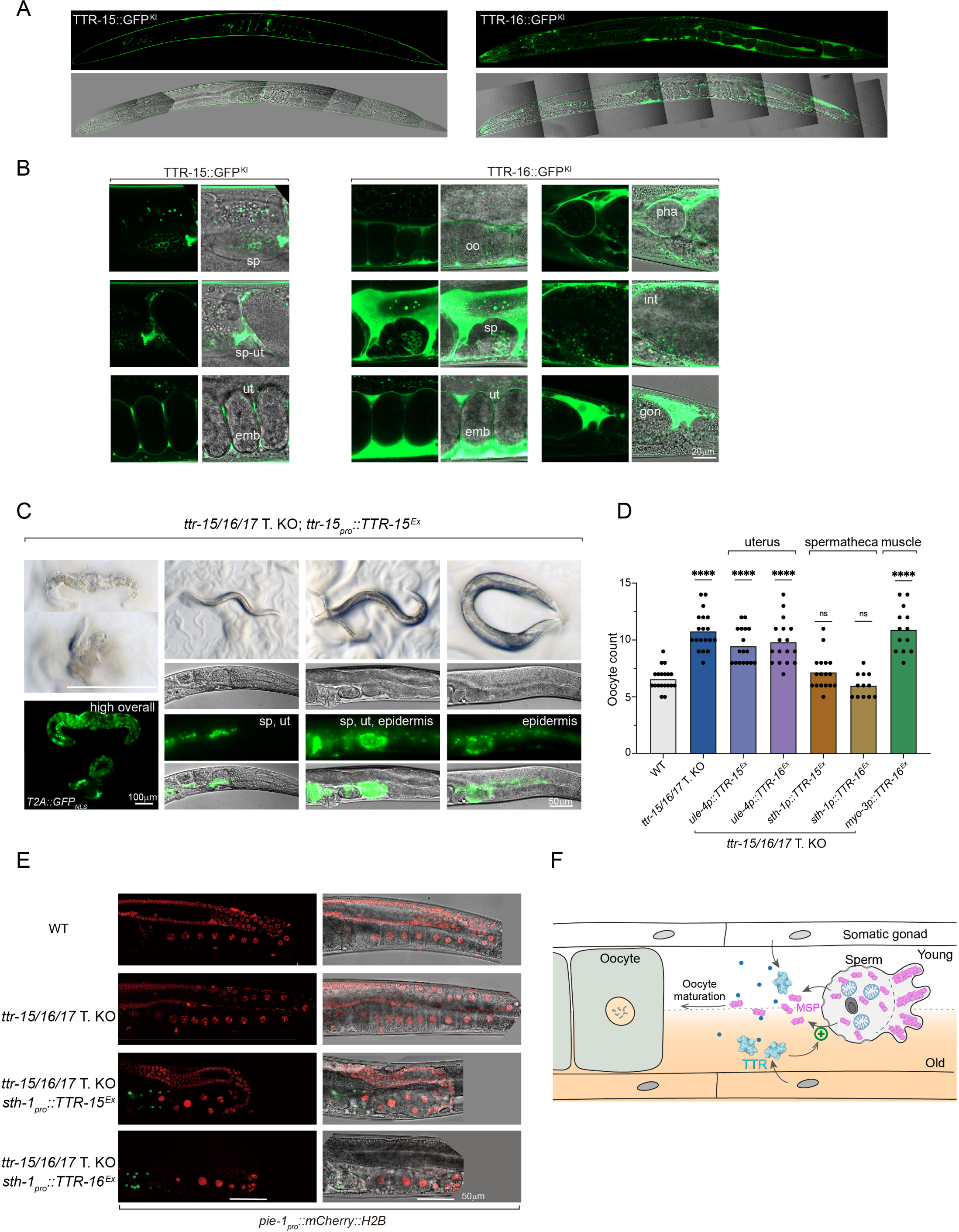
TTR-15 and TTR-16 Secretion in the Spermatheca Plays a Significant Role in Reproduction. A) Overview of TTR-15 and TTR-16 tissue expression and localization based on endogenously tagged GFP. TTR-15 is present inside the spermatheca and uterus. TTR-16 is present in: pseudocoelom; interspace between gonad sheath and germ cells; lumen of oviduct, spermatheca, and uterus. Refer to the supplemental figure S4 for T2A::GFP-NLS knock-in lines to label which cells express TTR-15, TTR-16, and TTR-17. B) Detailed view of TTR-15 and TTR-16 primary localization. Note that both TTR-15 and TTR-16 are secreted into the lumen of the spermatheca and uterus, surrounding the sperm. Abbreviations: sp, spermatheca; ut, uterus; sp-ut, spermatheca-uterine valve; emb, embryo; oo, oocyte; pha, pharynx; int, intestine. C) Overexpression of ttr-15 minigene carrying endogenous promoter and ORF leads to severe defects, including lethality. The expression of TTR-15 is labeled by T2A::GFP-NLS. The images show the F1 generation from transgene microinjection. Some larvae are lethal; those that grow to adults are nearly sterile. When TTR-15 is highly expressed overall, the animals show lethality and tissue disintegration (first column). When expression is prominent only in the spermatheca (sp) and uterus (ut), the animals are scrawny with severely deformed reproductive systems (second column). When expression is relatively broad, the animals show deformed gonads and sterility (third column). When expression is relatively enriched in the epidermis, the animals develop an abnormal body morphology and are rollers. D) The ovulation defects in triple KO animals are rescued by tissue-specific expression of TTR-15 or TTR-16. The graph quantifies the number of oocytes in one arm of the gonad. The expression of ttr-15 or ttr-16 in the spermatheca completely rescued ovulation, while the expression of ttr-16 in the muscle did not have rescuing effects. The expression of TTR-15 in the uterus showed some rescuing effect but was still far worse than the wild type. Refer to the supplemental figure S4 for rescuing by single copy insertion transgene. E) Overexpression of ttr-15 or ttr-16 in the spermatheca causes dysmorphogenesis of the gonad and aberrant oocyte development. In transgenic animals, the oocytes are irregularly packed in the oviduct, and the oocytes show decondensation of chromatin, a sign of disrupted oocyte arrest, which was not present in normal oocytes in wild-type animals. F) Proposed molecular action of TTR-15/16/17. After being secreted from the somatic gonad, TTR-15/16/17 plays a role in sustaining the release of major sperm protein (MSP) signals from activated sperm, particularly during the late phase of the reproductive period. MSP signals are crucial for inducing oocyte maturation. TTR-15/16/17 may interact with specific lipophilic signals in the extracellular environment to facilitate this process.

These results suggest that TTR-15 and TTR-16 are secreted extracellularly. To determine the specific cell types in which these genes are expressed, we generated expression reporter lines with T2A::GFP(NLS) knocked in at each gene’s C-terminus (**Figure S1C, S4A**). Analysis of GFP(NLS)-labeled nuclei indicated that TTR-15 is expressed in the spermatheca and uterus, while TTR-16 is expressed in muscles, neurons, and other tissues. TTR-17 displayed a similar expression pattern to TTR-16, although with much weaker intensity. Given that TTR-16 and TTR-17 are adjacent on the chromosome, their similar expression patterns suggest that one is likely a recent duplication of the other.

### Spermatheca-secreted TTR-15 and TTR-16 to Sustain Sperm Integrity

A single-copy transgene of P*ttr-15*::TTR-15 or P*ttr-16*::TTR-16 rescues the late-onset oocyte maturation defects in triple KO mutants (**Figure S4C**), supporting their roles in regulating reproduction. However, multicopy array transgenes of P*ttr-15*::TTR-15, with higher expression levels, cause severe developmental defects among F1 progeny (**Figure 5C**), with only one transmittable line out of 140 F1s. Abnormalities in F1 include dead embryos and young larvae, deformed reproductive systems, sterility, and roller phenotypes. In contrast, animals tolerate high levels of TTR-16, as similar multicopy arrays of P*ttr-16*::TTR-16 did not cause obvious growth defects in F1s.

Ectopic expression of TTR-15 and TTR-16 driven by the ubiquitous eft-3 promoter results in frequent embryonic and larval lethality, while epidermis-expressed TTR-15 (driven by the *dpy-7* promoter) consistently causes dumpy and larval arrest (not shown).

Since TTR-15 and TTR-16 proteins are secreted by somatic cells of the gonads and present in the luminal space, we tested gonad-specific expression driven by heterologous promoters (**Figure 5D**; **Figure S4D, E**). Spermatheca-expressed TTR-15 or TTR-16 (driven by the *sth-1* promoter) effectively rescues ovulation defects in triple KO mutants, while uterus-expressed TTR-15 or TTR-16 (driven by the *ule-4* promoter) provides partial rescue. Muscle-expressed TTR-16 (driven by the *myo-3* promoter) has no rescuing effect. These results suggest that the spermatheca and uterus are functional sites for TTR-15 and TTR-16 in regulating sperm activity, consistent with where endogenous proteins are detected.

In a subset of F1 animals with spermatheca-expressing TTR-15 or TTR-16 transgenes (**Figure 5E**), gonads exhibited dysmorphogenesis, including shortened and smaller distal gonads, erroneous turning, extra twists, bloated proximal oviducts, and thinned spermathecae. These nearly sterile animals were excluded from oocyte maturation analysis. Imaging mCherry::H2B-labeled chromosomes revealed diffused mCherry patterns in multiple oocytes within the oviduct (**Figure 5E**), indicating that these oocytes were no longer arrested in the diakinesis stage of prophase I of meiosis. This suggests excessive oocyte maturation induced by high-level expression of TTR-15 and TTR-16 in the spermatheca. Similar defects were not observed in F1 individuals with uterus-expressing TTR-15 or TTR-16 transgenes, likely due to the uterus being a more distant releasing site. These findings strongly support that TTR-15 and TTR-16 confer essential regulatory functions to reproductive processes (**Figure 5F**).

## Discussion

### Potential Roles of *C. elegans* TTR in Lipophilic Molecule Transport

Transthyretin, identified in the 1940s for binding thyroid hormones and retinol, is primarily associated with vertebrates^31,39^. In the genomic era, transthyretins were found to belong to a larger family called transthyretin-like proteins (TLPs) or transthyretin-related proteins (TRPs), present in various life forms, including bacteria and plants^40^. TLPs differ from transthyretin as they generally lack secretory signals, do not bind thyroid hormones, and function as enzymes by hydrolyzing 5-hydroxyisourate, an oxidative product of uric acid.

The genome of *C. elegans* contains two TLP orthologs and a large expanded family of 59 proteins with weak similarity to transthyretin or TLPs, designated as transthyretin-related proteins (TTRs)^38^. TTR-52 is well-characterized for its role in engulfing apoptotic cells by bridging phosphatidylserine (PS) on the dying cell surface and the phagocyte receptor CED-1^41^. Structurally, TTR-52 forms a dimer, unlike the transthyretin tetramer, and its open β-barrel-like structure resembles the C2 domain of protein kinase C, which binds PS^42^. Thyroid hormones have not been found in *C. elegans*^43^, suggesting TTRs in *C. elegans* are functionally distinct from TLPs and transthyretin. The evolutionary pathway of these proteins remains unclear, and the expression and function of most TTR family genes are largely unexplored.

Although *C. elegans* TTRs are unlikely to bind thyroid hormones, they may act as transport proteins for lipophilic molecules. Many transport proteins adopt a similar β-barrel-like structure^44,45^, such as fatty acid-binding proteins (FABPs), the lipocalin family including retinol-binding protein 4 (RBP4), NPC2 (Niemann-Pick C2) protein for cholesterol binding, and sex hormone-binding globulin. The abundance and location of such carrier proteins play a critical role in dictating the local availability of these molecules^46,47^. An example is the secretion of androgen-binding protein by Sertoli cells, maintaining high local concentrations of androgens crucial for spermatogenesis^48^. Future investigation is needed to determine if TTRs in *C. elegans* provide a similar function, given their significant role in regulating developmental timing and potentially reproductive aging.

### Critical Roles of Somatic Gonads in Sperm Maturation and Maintenance

The molecular regulation of sperm maturation is largely elusive for most animal species. In mammals, newly formed spermatozoa require additional processes to acquire the ability to fertilize^28,29^. Animal studies demonstrate that sperms released in the seminiferous tubules must pass through the epididymis to gain forward motility and the potential to interact with oocytes efficiently. Furthermore, freshly ejaculated human sperm are not immediately capable of fertilization; they need a period in the female reproductive tract to become competent. Human semen plasma contains over 2,000 proteins as well as other non-protein factors secreted from the testis, epididymis, and other accessory sex glands^49,50^, yet the function and significance of most of these substances in reproduction are not well understood.

In *C. elegans*, the spermatheca, the sperm ‘tract’ in hermaphrodites, plays an instructive role in sperm maturation^23^. Spermatids must enter the spermatheca to be remodeled and gain motility, though the molecular trigger for this activation is unknown. It is also unclear what roles the spermatheca plays in maintaining mature sperm. A commonly held but unexamined idea is that activated sperm remain autonomously competent for at least 3-4 days. Therefore, the observed decline in sperm quality in middle-aged TTR-15, TTR-16, and TTR-17 triple knockout animals was unexpected. To our knowledge, this is the first report of genetic mutants suggesting it is crucial for fully activated sperm to be maintained in the spermatheca throughout their reproductive lifespan.

The spermatheca may influence sperm at multiple levels, potentially involving the extracellular matrix, ion composition, energy resources, and specific maturation or maintenance signals. The specific roles of TTR-15, TTR-16, and TTR-17 in regulating sperm quality are currently unclear. These TTRs may affect sperm properties during spermatogenesis in the oviduct or maintain spermatozoa physiology in the spermatheca, or both. Notably, the spermatheca is the major expression site for these TTRs, which are secreted into its lumen where sperm are located. Understanding the molecular function of TTRs will provide insights into the interactions between the spermatheca and sperm in the context of sperm development and aging.

### Regulation of MSP Release via Unconventional Vesicles Remains Unclear

*C. elegans* sperm lack the ER-Golgi secretory apparatus, and thus conventional secretion pathways^15,18^. MSP lacks a secretory leader peptide and is released via double-membraned vesicles, as revealed by immuno-EM studies^21^. These studies suggest that MSP is localized in the interspaces between the outer and inner membranes of the vesicles. It is speculated that these vesicles may form from protruding cytoplasmic sacs driven by MSP polymerization, which extend, curve back to the cell membrane, and pinch off as double-membrane structures. The exact budding pathway of these MSP-containing vesicles remains experimentally uncharacterized.

MSP is released by hermaphrodite spermatids but not by male spermatids, implying that an unidentified signal in hermaphrodites triggers MSP release. The accelerated depletion of cytoplasmic MSP in TTR-15, TTR-16, and TTR-17 triple mutants supports that MSP release is regulated. It is conceivable that extracellular signaling coordinates with intracellular effectors to regulate the packaging of MSP into vesicles and the frequency of budding. These are critical questions for future investigation.

Over the past 20 years, an increasing number of both soluble cargos and integral membrane proteins have been found to reach the plasma membrane via unconventional pathways^51^. For example, FGF2, a leaderless cytoplasmic protein, translocates through the plasma membrane by forming a transport pore. Other proteins, like leaderless yeast Acb1, rely on membrane-bound organelles that divert from their normal function to become secretory. These pathways are diverse and heterogeneous, and a unified understanding of unconventional secretion is still lacking. MSP’s release in double-membraned vesicles is unique and has strong potential to provide insights into why and how cells utilize unconventional protein release mechanisms.

## Methods

### Caenorhabditis elegans Strains and Culture

The wild-type C. elegans strain used was Bristol N2. Strains were cultured on nematode growth medium (NGM) plates seeded with OP50 Escherichia coli cells as previously described^52^. The strains were maintained at 20°C, except where otherwise noted. Several temperature-sensitive infertile mutants were used as follows: spe-9(hc88) sperm are motile but defective in fertilization, fer-6(hc23) sperm cannot be properly activated and are therefore immotile, and fog-2(q71) is defective in sperm production. These mutants are fertile at 15°C and therefore maintained at this temperature. When used in experiments, they are transferred to 25°C from the point when oogenesis starts.

### Design and Construction of sgRNA for CRISPR-Cas9 System

We utilized the CRISPR-Cas9 system to design and construct sgRNA expression vectors. The Cas9 expression cassette was deleted from the original pDD162 (Peft-3::Cas9, U6 promoter::sgRNA) plasmid, resulting in a new plasmid with only the U6 expression cassette (designated as pQA1760). This served as the backbone vector for subsequent sgRNA expression vectors. sgRNA target sequences were selected using the CRISPR design tool (http://crispr.mit.edu). DNA oligos were designed based on the sgRNA targets, and the sgRNA vectors were generated using PCR-based cloning methods. High-fidelity enzymes, forward primer (5’-N20 GTTTTAGAGCTAGAAATAGCAAGTTAAAATAAG-3’), and reverse primer (5’-N*20 CAAGACATCGCAATAGGA-3’) were used. The N20 in the forward primer represents the 20 nucleotides gRNA target sequence, while the N*20 in the reverse primer is the reverse complement of the gRNA target sequence. After treating the PCR products with DpnI, transformation was performed using DH5α competent cells.

### Generation of Gene Deletion Mutants

We generated four gene deletion alleles, ttr-15(ybq209), ttr-16(ybq210), ttr-16(ybq246), and ttr-16 ttr-17(ybq211) mutants using CRISPR-Cas9 editing. Two sgRNAs were used for each gene: one targeting the first exon and the other targeting the last exon. The sgRNA-expression vectors were microinjected into the Cas9-expressing strain EG9887: oxTi1128 [mex-5p::Cas9(+smu-2 introns)::tbb-2 3’UTR + hsp-16.41p::Cre::tbb-2 3’UTR + myo-2p::2xNLS::cyOFP::let-858 3’UTR + lox2272] I.

#### Generation of ttr-15(ybq209)

The sgRNA targets are: sgRNA1 (AAGCAGACTATCACTGTCAA) and sgRNA2 (GACGACGAGATCACCTCAATC). F1 animals were singled, and F2 were screened using PCR with the forward primer (GCTTTGGCTTTTGTTGAGGC) and the reverse primer (AAATGTGGGTTCTGCAGCGA). The deletion allele was identified based on a band of 754

##### bp. ttr-15(WT, 1378bp)

**ATG**CGTGCTTTACTATTCACCTCTGTTGTTCTTTTGGCTTTGGCTTTTGTTGAGGCA AAGAAGCAGACTATCACTGTCAAG……GACGACGAGATCACCTCAATCTCTCCATACCTCATAATCACCCACAACTGCAACGTGAAGAAGGCCGGATGCAAGCGTGTTTCAGAGTATTTGATTCCAAAGGAGAAGATCGGTGGAACCTATGATATGACATACGTC ACTCTTGATATTCTTTCCGCTAAAGACAAGGAGAAGTGC**TAA**

##### ttr-15(ybq209)

ATGCGTGCTTTACTATTCACCTCTGTTGTTCTTTTGGCTTTGGCTTTTGTTGAGGCAAAGAAGCA-(1151 bp deletion)- ATCTCTCCATACCTCATAATCACCCACAACTGCAACGTGAAGAAGGCCGGATGCA AGCGTGTTTCAGAGTATTTGATTCCAAAGGAGAAGATCGGTGGAACCTATGATAT GACATACGTCACTCTTGATATTCTTTCCGCTAAAGACAAGGAGAAGTGC**TAA**

#### Generation of ttr-16(ybq210)

The vectors expressing sgRNA1 (CTTGTGCTCTTGAATGCACA) and sgRNA2 (CCTACGTCACCCTCGACATC) were microinjected into the EG9881 strain line. F1 animals were singled and screened using PCR with the forward primer (ATGTTGTCTCATGGCTGGCT) and the reverse primer (ATGCTCTTCAGGTCCTCGTTG), identifying mutant bands of 350 bp..

##### ttr-16(WT, 493bp)

**ATG**CGCTCGCTCGTCGTGTGTCTCTTGCTCGCCGCTTGTGCTCTTGAATGCA CAGCCCGTCTTCAGAATGTTA……TACGACATGACCTACGTCACCCTCGACATCA AGGTTCACGGAGAGAAGGAAAAATGCCAG**TAA**aaagtgcaaacttcctggattttattgactat

##### ttr-16(ybq210)

**ATG**CGCTCGCTCGTCGTGTGTCTCTTGCTCGCTTGATTCTATTGGTTGATTG AAGGGGGG-(463 bp deletion)-agtgcaaacttcctggattttattgactat

#### Generation of ttr-16(ybq246) under GFP::NKB-2 (ibp96, knock-in) background

Because ttr-16 and nkb-2 are close on chromosome, we generated additional ttr-16 deletion allele under the GFP::nkb-2(ibp96) background. The sgRNA1 (CTTGTGCTCTTGAATGCACA) and sgRNA2 (CCTACGTCACCCTCGACATC) vectors were microinjected into the LMW467(gfp::nkb-2) strain. F1 animals were screened using PCR with the forward primer (ATGTTGTCTCATGGCTGGCT) and the reverse primer (ATGCTCTTCAGGTCCTCGTTG), identifying mutant bands of 368 bp.

##### ttr-16(WT, 493bp) with GFP::NKB-2

**ATG**CGCTCGCTCGTCGTGTGTCTCTTGCTCGCCGCTTGTGCTCTTGAATGCA CAGCCCGTCTTCAGAATGTTA……TACGACATGACCTACGTCACCCTCGACATCA AGGTTCACGGAGAGAAGGAAAAATGCCAG**TAA**

##### *ttr-16(ybq246)* with GFP::NKB-2

**ATG**CGCTCGCTCGTCGTGTGTCTCTTGCTCGCCGCTTG-(417bp deletion)- ACATCAAGGTTCACGGAGAGAAGGAAAAATGCCAG**TAA**

#### Generation of ttr-16 ttr-17(ybq211)

The sgRNA1 (GGCTCAAATTCATACCAACA) and sgRNA2 (ACGTCACCCTCGACATCAAG) vectors were microinjected into the EG9881 strain line. F1 animals were screened using PCR with the forward primer (TTTCAGCCATGTCTCGCCTT) and the reverse primer (ACTCCTCGATTTCCAACGGG), identifying mutant bands of 465 bp.

##### ttr-16 ttr-17(WT, 3403bp)

**ATG**TCTCGCCTTGCTATCCTTGTTCTTTTGTCACTCGCTGTATTTGAAGTATC CGCCAAACTTCAAAATGTAACTGTAAAAGGAATTGCCGTTTGTAACAAGAGAAGAATGGCTAATTCCCATGTTATTCTGATTGATAAGGATACCTgtaagtgtgacaaggaaaaatattaatattattccataagtttcagTGGACCCAAATGACGAATTGGCTCAAATTCATACCAACAAGGAAGGAGAATTCGAGTTG……TGGAGAGAAGACCATCTGCAAC**TAA**ttatgatttttttgtttca actcatataattgta……**ATG**CGCTCGCTCGTCGTGTGTCTCTTGCTCGCCGCTTGTGCTCTT GAATGCACAGCCCGTCTTCAGAATGTTA……TACGACATGACCTACGTCACCCTC GACATCAAGGTTCACGGAGAGAAGGAAAAATGCCAG**TAA**

##### ttr-16 ttr-17(ybq211)

**ATG**TCTCGCCTTGCTATCCTTGTTCTTTTGTCACTCGCTGTATTTGAAGTATC CGCCAAACTTCAAAATGTAACTGTAAAAGGAATTGCCGTTTGTAACAAGAGAAGAATGGCTAATTCCCATGTTATTCTGATTGATAAGGATACCTgtaagtgtgacaaggaaaaatattaatattattccataagtttcagTGGACCCAAATGACGAATTGGCTCAAATTCATAC-(3130bp deletion)- ACCCTCGACATCAAGGTTCACGGAGAGAAGGAAAAATGCCAG**TAA**

#### Generation of Double and Triple Mutants by Crossing

The ttr-15(ybq209) was crossed with the ttr-16(ybq210) to obtain the double mutant ttr-15(ybq209); ttr-16(ybq210). The ttr-15(ybq209) was crossed with the ttr-16-17(ybq211) to obtain the triple mutant ttr-15(ybq209); ttr-16 ttr-17(ybq211).

### Generation of Endogenous GFP or T2A::GFP Tagging Alleles

We constructed three GFP inserted alleles: ttr-15::gfp(ybq203), ttr-16::gfp(ybq204), and ttr-17::gfp(ybq205). We also generated three T2A::GFP inserted alleles: ttr-15::T2A::gfp(ybq206), ttr-16::T2A::gfp(ybq207), and ttr-17::T2A::gfp(ybq208). The sgRNA target sequences were selected near the start codon of ttr-15, ttr-16, and ttr-17, choosing two closely spaced sgRNAs. For the homologous arms, approximately 1 kb upstream of the sgRNA1 target sequence and 1 kb downstream of the sgRNA2 target sequence were selected. Vector plasmids carrying GFP or T2A::GFP and homologous arms were constructed using homologous recombination, and the PAM site of the sgRNA was modified to a synonymous coding sequence to prevent repeated cleavage by Cas9. The Cbr-unc-119 expression cassette as a selection marker is embedded within the third intron of the GFP ORF. The repair template plasmid, sgRNA1, and sgRNA2 were co-injected into the gonads of the EG9881 strain. The injection concentration for the repair template plasmid was 40 ng/μl, and for the sgRNAs, it was 10 ng/μl. F1 animals with restored locomotion were selected and raised individually, and the locomotion of the F2 generation was examined as screening criteria. After confirming GFP fluorescence under a microscope, they were crossed three times with the wild-type N2 strain before use.

#### ttr-15::GFP(ybq203) *and* ttr-15::T2A::GFP(ybq206)

The sgRNA1 (GACGACGAGATCACCTCAATC) and sgRNA2 (agctcactgctcgagctgga) were used to generated the knock-in alleles.

CcGACGACGAGATCACCTCAATCTCTCCATACCTCATAATCACCCACAACTGCAACGTGAAGAAGGCCGGATGCAAGCGTGTTTCAGAGTATTTGATTCCAAAGGAGAAGATCGGTGGAACCTATGATATGACATACGTCACTCTTGATATTCTTTCCGCTAAAGACAAGGAGAAGTGC- GFP or T2A::GFP insert - **TAA**gaaaatgttttttttgtttgtttgcttgtttggaagggaaggactttctatctcttttaattcaacaataaactattggaaaaccttgaaatt ttaaccttgaactgtaagaaaagttgcgtgattatgttgacaattttgccaagtatatctttgtggatatcacaataaacgaagtcaaagcacgaaatattacggaaacacaaaattaatgagaatgcgcaacatatttgaccgcaaaatatctcgtagcgaaaactacagtaattcttcaaaa gactactgtagcgcttgtgtcgatttacgagctcgatttttgaaatgaatcagactagaagaaaaggaggaaaatattgaacatcaattgaacatcaattcaaaaagtcgaacccttgactacagtagtcttctaaagaattactgtagttttcgctacgagatattttgcgcgtcaaatatgttgcgcaatacgcatcctcagaattgtgtgttctcgtaatgtcttgaaaattttccatttcaacatcaaataagcaaatctaaaaatgtgggttctgcagcgaccactatgactgtgatcgtggcaagacccactcagaaaactacgtgttcctttaaacaatacatttttaagtattgtaggtataaaaattgttggctagcagtctaggctgcctttttcagtcgacaaacttctaatttaatcggcgggtcttcaaaaagtcgtttctttgaaaatataaagctttatatatttatatattaaaaattttgattacatgatatcaaaagcgaccagtttgcataaaaattatcaaccaacaatatcaatatcttcttctacctcctgatcctcctccttcacaatgctcagctcactgctcgagctggaagCagtcgacggga

#### ttr-16::GFP(ybq204) *and* ttr-16::T2A::GFP(ybq207)

The sgRNA1 (TACGTCACCCTCGACATCAA) and sgRNA2 (aacgaggacctgaagagact) were used to generated the knock-in alleles.

CTACGACATGACATACGTCACCCTCGACATCAAAGTTCACGGAGAGAAGGAAAAATGCCAG-GFP or T2A::GFP insert - **TAA**aaagtgcaaacttcctggattttattgactatctaaatatatattttttctatatgatttttctcaattttccattaataaaaacatgaaataa atgtttagttttttcaacgaggacctgaagagactagCgatcaa

#### ttr-17::GFP(ybq205) *and* ttr-17::T2A::GFP(ybq208)

The sgRNA1 (GCTCAAATTCATACCAACAA) and sgRNA2 (ctgccaactgacgaaagaga) were used to generate the knock-in alleles.

TTaGCTCAAATTCATACtAACAAcGAAGGAGAATTCGAGTTGTTCGGAGAGGAAGATGAGATTGGCAAAATTGAGCCATACATTCGTATTCATCACAGTTGCAATACCAAACCAgtaagtatttctaatttaatctaatcttatgagtgaaactgtaggaataaattagaaaatctggctggtaataaggaaaatggcaatttgaaaaagtttgatattttcttgaattattatacattattataaattatcatgaataatacattgatgagaagcagattataaaggtgcatatatagatacaaattaaccttgtgaattgaaaatatctagtacaatatgacacttaaataaccctcccaagtaggcaaatatctcttcctgactgaaagccaatttacgctctcctgatgataaattgaaacttttggtaaatttattgcaaatcagaaatacgcaaatctacacagtgcaaatactgagcgtggtttgaaatgcaaaatataggcggtaggtatttaggcacgtaaacatgcacacctacttagcaatagctacttcgaatataaatttgaaaaacttaattgaagaaatttaaaaaatgttatgtttccagGGATGTGAACGCGTTTCCGAATATCAAAT TCCACAAGAAAAAATCGGAGAGGTCTATGACATGACTTATGTTACCCTTGATATTATTGTTCATGGAGAGAAGACCATCTGCAAC- GFP or T2A::GFP insert - TAAttatgatttttttgtttcaactcatataattgtatgtacataaattcgtattttaccctaaaacgttttgaactatttctggtttttgtttttaattgattcggagtacaacAAGctgcTaactgacgaaagagaaac

#### DNA Sequence for GFP carrying Cbr-unc-119 expression cassette

**atg**agcaaaggagaagaacttttcactggagttgtcccaattcttgttgaattagatggtgatgttaatgggcacaaattttctgtc agtggagagggtgaaggtgatgcaacatacggaaaacttacccttaaatttatttgcactactggaaaactacctgttccatgggtaagtttaaacatatatatactaactaaccctgattatttaaattttcagccaacacttgtcactactttctgttatggtgttcaatgcttctcgagatacccagatcatatgaaacggcatgactttttcaagagtgccatgcccgaaggttatgtacaggaaagaactatatttttcaaagatgacgggaactacaagacacgtaagtttaaacagttcggtactaactaaccatacatatttaaattttcaggtgctgaagtcaagtttgaaggtgatacccttgttaatagaatcgagttaaaaggtattgattttaaagaagatggaaacattcttggacacaaattggaatacaactataactcacacaatgtatacatcatggcagacaaacaaaagaatggaatcaaagttgtaagtttaaacatgattttactaacataacttcgtataatgtatgctatacgaagttatttctagacattctctaatgaaaaaatctttcagttgaaattgaaaatgagttaaagttggagtttttattgaaaacagatttccgtgtgattagtgtttttagcgagtgtgacaggacagcgaaaaaatatagaaacaaggggggaactgaaaagcttaggaatgcattgaacatgagaaggggaaggggaaggaacaaactagacaggaattattggaatttaatcacatttggagttttttttctattcgacagaataattatccagaacatttttgtattaaatatTTATGCATCATATGAGTAGTCGGCTTTGTTGTGCATGACGAGTTTGTTATCGACGAAATAGAAGCTGTCAGAACGAGTCTCGTTTGGATTGTTGATCATGTCGTCCActgaaaaagagattagtcttttgaattgtactttttagaataatgactcacTGAGTTGTTGAGAGAGTTGAGGGAACTCATAGATATGTTCACAGTTGTTTCGTGAATTCGGAATACAGAATCCGAATTCAAAGTCAAAACACTTCAGAAGGCGATCTTTGAAGAAGTGACGTTCGATCATTCGGAAATGATGGATTGGGATGTCTCCTACTTTGAATTCCACAGTTGCTCCGACCGTTTTGAGTTTCAAAAAGTTTGGAGCAAATCTATAACGCACGTATCTTGCCGACTCCTGTGGCGACTCATCATTCTCCTGATCGTTCTCCGGTTTGGCGATCTCGAAAAGCACTTGCTCAGTGTCCAGATCACGGATTTGGAACTTGGTGAACTCGATGTTATAGATGTTCGCAGATGGGGAGCATAAGAATcctaaatttatgttttaaactgaaatccaaagggagcaagataCCTTGAGTGATTCCCGGAAGTGCTAAAACGTCGTTCGGAGTGATTTGAGCTTTCTTCGCAAGCTCCGATTCCGTTGTGATTCCTTGTTCGGTGCTTGGTGGTGGCCGTGGCATC TGgaaatatggaaaagttcaacaaaaagaaaagagaaaagaatgaaatcggatatcaagagttagttgagcggtttctctagttttctgagtctcacCTGCGACGGGAAGGTCGCCGAGCCGGGTGGAATCGATTGTTGTTGCTCGGCTTTCATatcggtttggttggaagcggctgaaaacggaaagaagtggaagaaggaaaagagtgtggtgtgacaggaaaatggtaattagagggtgccaaataaccagctatattttgtttttttttgaaaacatttttaaaaagaaaaatacgataatgatatcagatggatttccggaaaactggtatgaaaaatttcaacctttttgagtacatgtaatcaaaatacactttgtaaattatcatttttattgaaactccaccatttttctatttataacgctaataatttgaaaaagaaacctgttgcgaaccgcggggtgaatcccaaaaacgaatgcgttttggtggagtgattgattcgaatcgaagaagaaaaagaagaagacgtggaatagagagctcactcttaaccgagcagcacacaccgacagaaaaaaaaatgaaatgaatgagggtcttcttcttcttcttcttcgaatgattgacagaaatgggaaaaagaggaagattgagaagggaaaaaggaaggagaaaagaagcagaagaagacgtcagagaggagaggaacgagcggaaaagcagcgggcgcaagtcatagaagtagcagagctggggagaagaagacactatccaagaaaggaatgacgagagagtatgcaaaggggtatagggtgcagacagaataggaacagaataacagatgatgagccaagaagagttgaaaagggcgatgaatttgtcatgtaacttaatttgggtcaatttgagcatgatgaattgaaatcatcccttgttgggagttaataaccggtttgttatcagaaaccctgtaatagaagggcgccctaactttgagccaattcatcccggtttctgtcaaatatatcaaaaagtggtcaactgacaaattgtttttgatattataataaacattttatccgttaacaattttcgaatactttttacaaggacttggataaattggctcaaagataacttcgtataatgtatgctatacgaagttattaactaatctgatttaaattttcagaacttcaaaattagacacaacattgaagatggaagcgttcaactagcagaccattatcaacaaaatactccaattggcgatggccctgtccttttaccagacaaccattacctgtccacacaatctgccctttcgaaagatcccaacgaaaagagagaccacatggtccttcttgagtttgtaacagctgctgggattacacatggcatggatgaactatacaaa

#### DNA sequence for T2A

GAGGGACGTGGATCCCTTCTTACCTGCGGAGACGTCGAGGAGAACCCAGGACCAATGGCAAGTTTGTACAAAAAAGCAGGCTCGCCAAAGAAGAAGCGTAAGGT G

### Generation of Single-Copy Integrated Transgene

We used the miniMos technology to construct two strains: Pttr-15::TTR-15::T2A::GFP(ybqSi280) and Pttr-16::TTR-16::T2A::GFP(ybqSi281). Using the pCFJ797 plasmid as a backbone, we constructed the Pttr-15-ttr-15-T2A-gfp (pQA2193) and Pttr-16-ttr-16-T2A-gfp (pQA2194) plasmids. The promoter region of ttr-15 contains the 1605 bp upstream of the start codon, and the promoter region of ttr-16 contains the 2080 bp upstream of the start codon. The injection mixture contained 10 ng/μl Prab-3::mCherry (pGH8), 1 ng/μl Pmyo-2::mCherry (pCFJ90), 50 ng/μl Peft-3-Mos1-transposase (pCFJ601), and 50 ng/μl miniMos plasmid carrying the target expression cassette. The mixture was injected into the gonads of N2 animals. Post-injection, P0 worms were placed in groups of three on each NGM plate and cultured at 25 °C. After 48 hours, 500 μl of a 25 mg/ml G418 solution was added, and after 7-10 days, healthy worms lacking the mCherry co-marker were selected from the plates. Once homozygosity was confirmed, the strains were crossed three times with N2 before use.

### Generation of Multicopy Array Transgene

For analyzing the expression pattern of TTR genes, we generated reporter constructs, such Pttr-15::TTR-15::T2A::GFP, and microinjected the plasmids into the gonads of the N2 strain. The injection mix consisted of 10 ng/μl of the target plasmid, 10 ng/μl Pmyo-3-mCherry, and 1 ng/μl myo-2-mcherry.

For rescuing experiments, plasmids Pule-4-ttr-15::T2A::gfp (pQA2195), Pule-4-ttr-16::T2A::gfp (pQA2196), Psth-1-ttr-15::T2A::gfp (pQA2199), Psth-1-ttr-16::T2A::gfp (pQA2200), and Pmyo-3::ttr-16::T2A::gfp (pQA2262) were constructed and injected into the strain ttr-15(ybq209); ttr-16 ttr-17(ybq211) (BLW2214). The injection mix contained 10 ng/μl of the respective target plasmid and 10 ng/μl Pmyo-3-mCherry.

### CRISPRi System for Gene Knockdown

The CRISPRi system can simultaneously knock down multiple genes, enabling the observation of resulting phenotypes in offspring. For each gene, two sgRNAs targeting the transcription start site (TSS) region are designed. To target 4-6 genes concurrently, the pool of these sgRNA-expressing plasmids was microinjected into the gonads of a single-copy insertion strain DZ89 of unc-119(0); Peft-3-dCas9(dPi)::GFP(dPi) unc-119(+) II. The injection mixture contained 10 ng/μL Pmyo-3::mCherry, 1 ng/μL Pmyo-2::mCherry, and 10 ng/μL sgRNA vector for each gRNA target. Following injection, the general development and morphology of F1 progeny were examined and scored under a fluorescent stereo microscope.

### Analyzing Overall Development of ttr-15, ttr-16, and ttr-17 Mutants

To compare the developmental rates between the ttr-15, ttr-16, and ttr-17 mutants and N2 wild-type worms, the animals were synchronized to young L1 stage via egg bleaching and raised on NGM plates with an OP50 bacterial lawn. After 36 hours of culture, most N2 animals had reached the L4 stage. Using the L4 stage as a reference, we counted the number of animals at specific stages: younger than L4, L4, and adult stage. The experiments were repeated three times on different dates. A pie chart was generated based on the counts to visualize the differences between the distinct genotypes.

### Analysis of Oocyte Maturation

The ttr-15, ttr-16, and ttr-17 mutants exhibited abnormalities in oocyte maturation. To analyze these abnormalities, we quantified the timing of the appearance of the stacked oocyte phenotype and the number of oocytes in the germ arm.

For the timing of defect onset, we assessed the presence of stacked oocytes in four stage-groups: 24, 48, 60, and 72 hours post-mid-L4. We observed and scored whether the arrangement of oocytes in the oviduct was normal or abnormally stacked for each individual under a microscope (Olympus MVX10, 40x magnification). We then calculated the percentage of animals with obvious oocyte stacking in each group.

To determine the number of oocytes present in the oviduct, we selected animals at 72 hours post-mid-L4 stage and counted the number of oocytes in one gonad arm, from the most proximal oocyte to the most distal one at the gonad turning point.

### Assays on Progeny Count

To count the number of progenies, single mid-L4 hermaphrodites were picked onto NGM plates with an OP50 bacterial lawn and allowed to grow and lay eggs. Each individual was transferred to a new plate with food every 24 hours. All hatched young larvae on the plates were counted. The counts from these three plates (0-72 hours post mid-L4) were added up to obtain the total number of progenies for each individual. A graph was generated from the counts of multiple individuals, with each data point representing the progeny count of one individual animal.

### Assessment of Embryonic Viability

It was observed that older ttr-15, ttr-16, and ttr-17 triple KO animals produced some dead eggs. To assess embryonic viability, we selected animals at 48 hours post-mid-L4 stage. Groups of 10 animals were placed on a single NGM plate with food and allowed to lay eggs for 24 hours. The parent adults were then removed from the plate, and the eggs were allowed to hatch for 12 hours. The number of hatched larvae and unhatched fertilized eggs were counted.

Unhatched fertilized eggs are morphologically distinctive as they are more round instead of oval-shaped and darker in color. These unhatched fertilized eggs can be differentiated from unfertilized oocytes, which appear bigger, rounder, and lighter in refraction due to the lack of divided cells in embryos.

### Crossing with Males to Rescue Ovulation

To evaluate the potential rescue of ovulation defects in ttr-15, ttr-16, and ttr-17 triple KO hermaphrodites, the following crossing procedure was conducted. Preparation of hermaphrodites: Wild-type and ttr-15, ttr-16, and ttr-17 triple KO hermaphrodites at the mid-L4 stage were selected and cultured on a fresh plate at 20°C for 48 hours to obtain Day-2 adults. Preparation of males: The him-5(ok1896), spe-9(hc88); him-5(ok1896), and fer-6(hc23); him-5(ok1896) strains were maintained at 15°C. Gravid adults were transferred to NGM plates and allowed to lay eggs at 25°C for 2 hours to obtain a relatively synchronous population, which was then cultured at 25°C. Males at the L3 stage were selected and transferred to a new plate, then cultured at 25°C for 24 hours to reach young adulthood and be ready for crossing. Crossing procedure: 30 Day-2 hermaphrodites and 10 young adult males were placed together on a plate for crossing at 25°C for 24 hours. After crossing, the hermaphrodites were transferred to a new plate and allowed to grow and lay eggs at 20°C for 24 hours. These hermaphrodites (96 hours post-mid-L4) were then imaged using an Olympus MVX10 microscope at 60x magnification and analyzed for phenotypes associated with oocyte maturation.

To assess the progeny number after crossing with males, the following procedure was implemented. Individual hermaphrodites at the L4 stage (either N2 or ttr-15/16/17 triple KO) were placed together with a single L4 male of the him-5(ok1896), spe-9(hc88); him-5(ok1896), or fer-6(hc23); him-5(ok1896) on a single plate. The crossing plates were kept at 25°C for 36 hours. After this period, each hermaphrodite was transferred to a new plate and allowed to continue laying eggs at 25°C. The hermaphrodites were transferred to a new plate every 24 hours. The total number of viable progenies produced by each hermaphrodite was counted on these plates.

### Whole-mount DAPI Staining

Animals of the mid-L4 stage were cultured for 0, 24, 48, 60, and 72 hours. They were then collected and rinsed in 1 mL of M9 buffer in 1.5 mL microcentrifuge tubes and centrifuged at 3000xg for 2 minutes. This washing step was repeated three times. Afterward, the animals were fixed in 200 µL of 2% paraformaldehyde (PFA) at room temperature for 15-30 minutes. The animals were then washed three times with 1 mL of PBST (1x PBS, 0.5% Triton X-100). The animals were then incubated in the PBST solution containing 10 ng/mL DAPI (4’,6-diamidino-2-phenylindole) at room temperature for 40 minutes. Post-staining, the samples were washed three times with the PBST solution, followed by one more wash with M9 buffer. The animals were then mounted and imaged on an Olympus MVX10 with 60x magnification, focusing on the spermatheca and oocyte maturation.

### Sperm Activation Assays

Males of N2 and ttr-15/16/17 triple KO mutant strains at the mid-L4 stage were placed on NGM plates seeded with OP50. After 48 hours, the males were dissected to release sperm into sperm buffer (50 mM HEPES, 25 mM KCl, 45 mM NaCl, 1 mM MgSO4, 5 mM CaCl2, 10 mM Dextrose; pH 7.8) with or without Pronase (200 μg/mL)^53^. The sperm were then incubated for 15 minutes and imaged to observe and compare sperm activation.

### Immunostaining of MSP

Hermaphrodites at 24, 48, 60, and 72 hours post-mid-L4 were prepared, dissected, and immunostained to analyze the protein level of MSP in vivo. For each stage, 200 worms were selected and washed in 1 mL of M9 buffer in a 1.5 mL centrifuge tube. After a 5-minute settling period, the worms were rinsed once with 1 mL of M9 buffer to remove residual bacteria. Next, 100-150 worms were transferred onto a glass slide containing 200 μL of PTW (1XPBS and 0.1% Tween 20) solution supplied with 0.2mM Levamisole. It took 1-2 minutes until the worms were fully paralyzed, and their gonads were then dissected out. The dissected gonads were transferred to a new 1.5 mL centrifuge tube and centrifuged at 8000 rpm for 2-3 seconds. Subsequently, the tissues were fixed in 200 μL of 4% PFA in PBS at room temperature for 30 minutes. The samples were washed twice with PTW solution. After washing off the PFA, 200 μL of ice-cold 100% ethanol was added, and the sample was incubated at −20°C for 10 minutes. The samples were then washed three times with 200 μL of PTW with additional 0.5% BSA, with each wash lasting 10 minutes. The samples were incubated in primary antibody solution (PTW with 1:100 diluted MSP antibody and 1% donkey serum) at room temperature for 2 hours. The samples were washed three times, each with 200 μL of PTW/BSA solution for 10 minutes. The samples were incubated in the secondary antibody solution (PTW with 1:1000 diluted Alexa Fluor 488 Donkey anti-Mouse IgG (H+L) and 1% donkey serum) at room temperature for 1 hour, followed by three washes with 200 μL of PTW/BSA solution, each lasting 10 minutes. Finally, the samples were sealed using a mounting agent and imaged under a ZEISS LSM 980 with Airyscan 2.

### Image Acquisition

For analyzing fluorescence-labeled sperm, we used the marker line ujIs113 [pie-1p::mCherry::H2B::pie-1 3’UTR + nhr-2p::his-24::mCherry::let-858 3’UTR + unc-119(+)]. Imaging of the complete spermatheca was performed using an Olympus MVX10 microscope at 60x magnification in Z-stack mode with a 1 μm step size. The sperm were visually identified and counted.

For imaging TTR-15, TTR-16, and TTR-17 GFP fusion or T2A::GFP knock-in lines, animals were immobilized between an agarose pad and a cover glass. Images were acquired on a ZEISS LSM 980 microscope with Airyscan 2 using a 63x (NA 1.4) lens. When needed, images were stitched together using Adobe Photoshop.

The immunostaining signals of MSP were captured on a ZEISS LSM 980 Airyscan 2 microscope at 63x (NA 1.4) using identical acquisition parameters, such as laser power and gain values. To measure the intensity, the fluorescent positive areas were selected by manually thresholding the images. The average signal intensity of each target image was then recorded. The intensity data were further analyzed for statistical significance using Prism GraphPad.

### Statistical Analysis

The differences between experimental and control groups were analyzed using a t-test. The data are presented as Mean ± SEM in a column graph with each data point shown. Statistical significance is indicated as follows: ns (not significant), * (p < 0.05), ** (p < 0.01), *** (p < 0.001), and **** (p < 0.0001).

## Acknowledgments

We express our gratitude to Long Miao and his lab members for providing strains and engaging in helpful discussions; Yan Zou, Xinjian Wang, and the Zou lab members for reagents and discussions; Andrew Singson for supplying strains; Qian Bian, Di Chen, and David Greenstein for their valuable discussions; and Dandan Yü for assistance with immunostaining. We thank the Molecular and Cell Biology Core Facility (MCBCF) and the Molecular Imaging Core Facility (MICF) at the School of Life Science and Technology, ShanghaiTech University for providing technical support. Some strains were provided by the CGC, which is funded by NIH Office of Research Infrastructure Programs (P40 OD010440). This work was supported by the lab start-up fund from ShanghaiTech University to Y.B.Q. and by National Science Foundation of China (Grant no. 32370601) to Y.B.Q.

---

**Figure S1.**
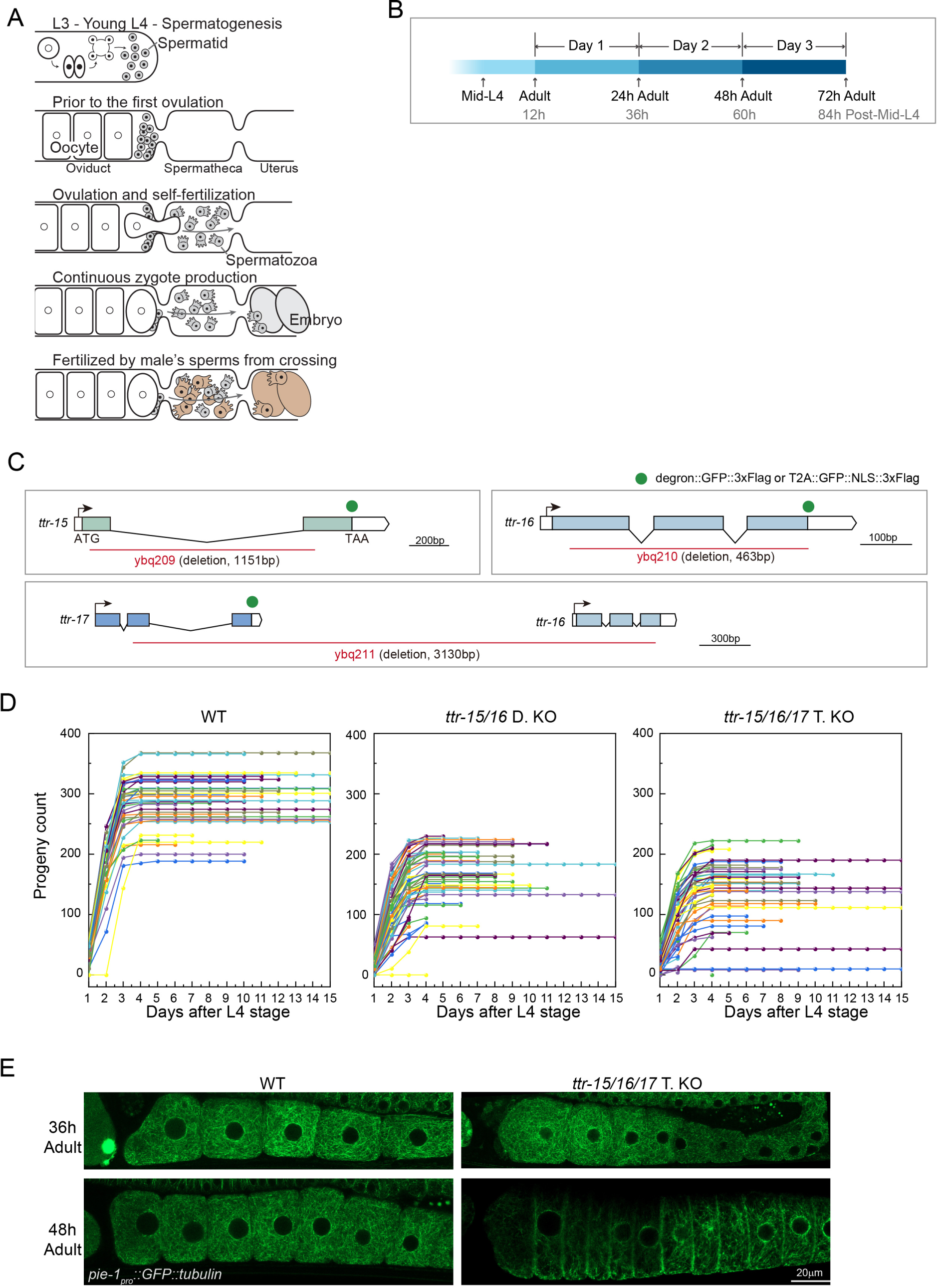
Oocyte Defects in TTR-15, TTR-16, and TTR-17 Triple KO Animals. A) The drawing illustrates the generation sequence of sperm and oocytes, ovulation, and self- and cross-fertilization. Spermatogenesis precedes oogenesis in hermaphrodites, giving rise to a limited number of self-sperm. Maturing oocytes move into the spermatheca, prompting spermatid activation into motile spermatozoa. Following fertilization, zygotes form and enter the uterus. Crosses with males bring male sperm into the uterus and spermatheca, with male sperm being more competent than self-sperm. B) Illustration of stage definitions in this study. The stages are defined as follows: Mid-L4 are visually assessed based on the morphology characteristic of the L4 larval stage. Adults are marked by the formation of zygotes. Young Adult are the stage between the L4-adult molting and the adult stage. For experimental convenience, “hours post-mid-L4” is used, which is translated to adult stages in the paper. C) Illustration of deletion mutants for ttr-15, ttr-16, and ttr-16 ttr-17. The GFP or T2A::GFP-NLS endogenous tag is inserted at the C-terminus of the gene before the stop codon. D) Progeny count by tracking individual animals. The indicated genotypes include wild type, ttr-15/16 double KO, and ttr-15/16/17 triple KO. Each colored line represents a single animal, with the end of the line indicating the demise of the individual. E) Imaging GFP-labeled microtubules in oocytes of wild-type and triple KO animals at the first and second days of adulthood. MSP triggers microtubule reorganization such that microtubules are evenly dispersed in the cytoplasm, as observed in wild-type and first-day triple KO animals. In the second day of triple KO animals, microtubules are cortically enriched, suggesting a lack of MSP signals.

**Figure S2.**
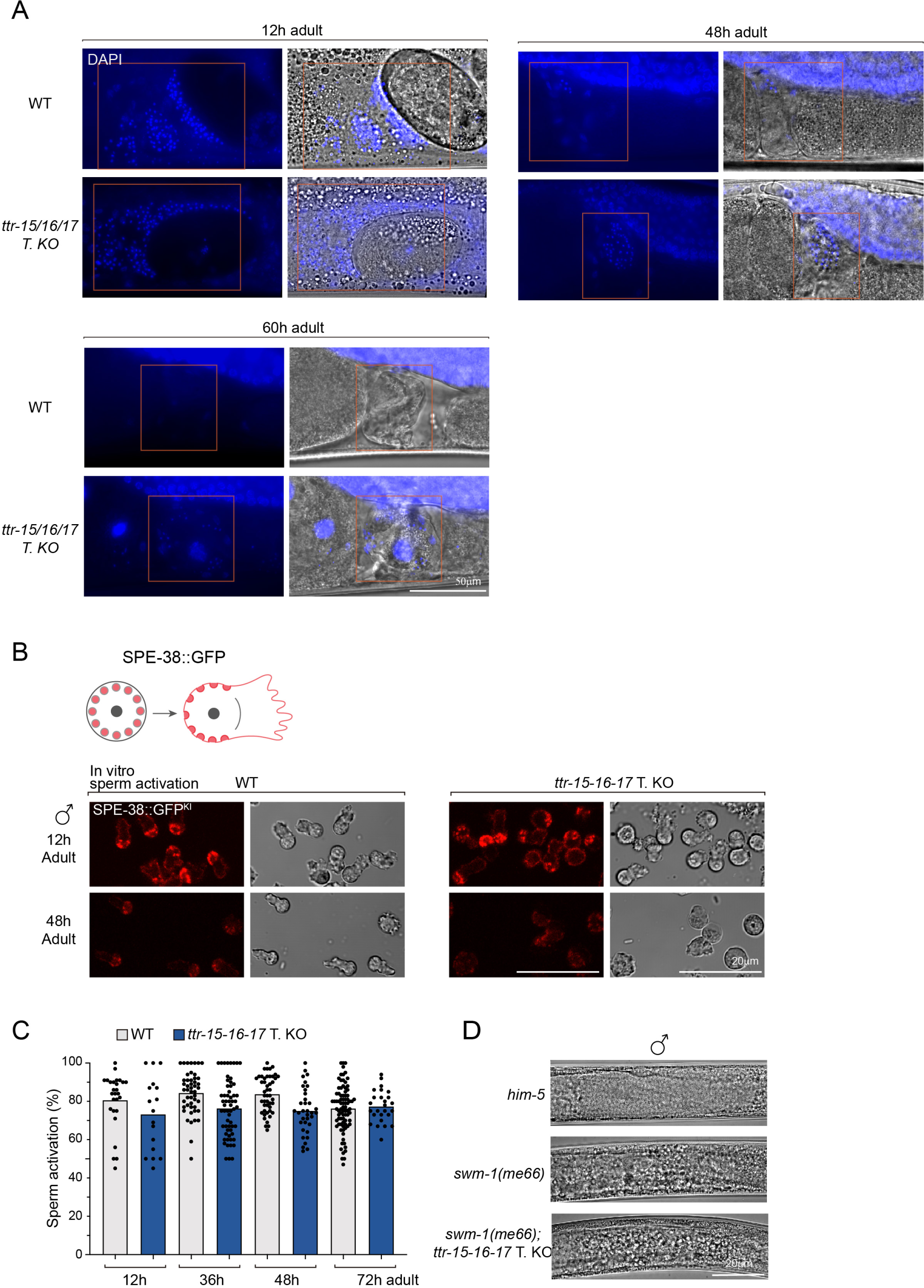
Persistent Sperm in Triple KO and Male Sperm Activation In Vitro. A) Whole mount DAPI staining of wild-type and triple KO animals to visualize sperm. The red-lined rectangle indicates the position of the spermatheca. DAPI-labeled sperm are present in older mutant adults, while sperm are depleted in most age-matched wild-type animals. B) In vitro male sperm activation is normal for triple KO. The pattern change in SPE-38::mCherry (knock-in) is used to label sperm activation. The presence of pseudopod extension under DIC microscopy is detected in triple KO sperm. C) Quantification of in vitro male sperm activation for wild-type and triple KO animals. Each data point represents the percentage of activated sperm based on morphology within one captured independent field under DIC microscopy. D) swm-1 mutant male sperm are prematurely activated in the male gonads. The male sperm of triple KO mutants are also prematurely activated in the absence of swm-1, suggesting that sperm activation in triple KO is normal in vivo. The image shows the male gonad, with sperm activation marked by the refractory change from smooth, immotile spermatids to grainy spermatozoa.

**Figure S3.**
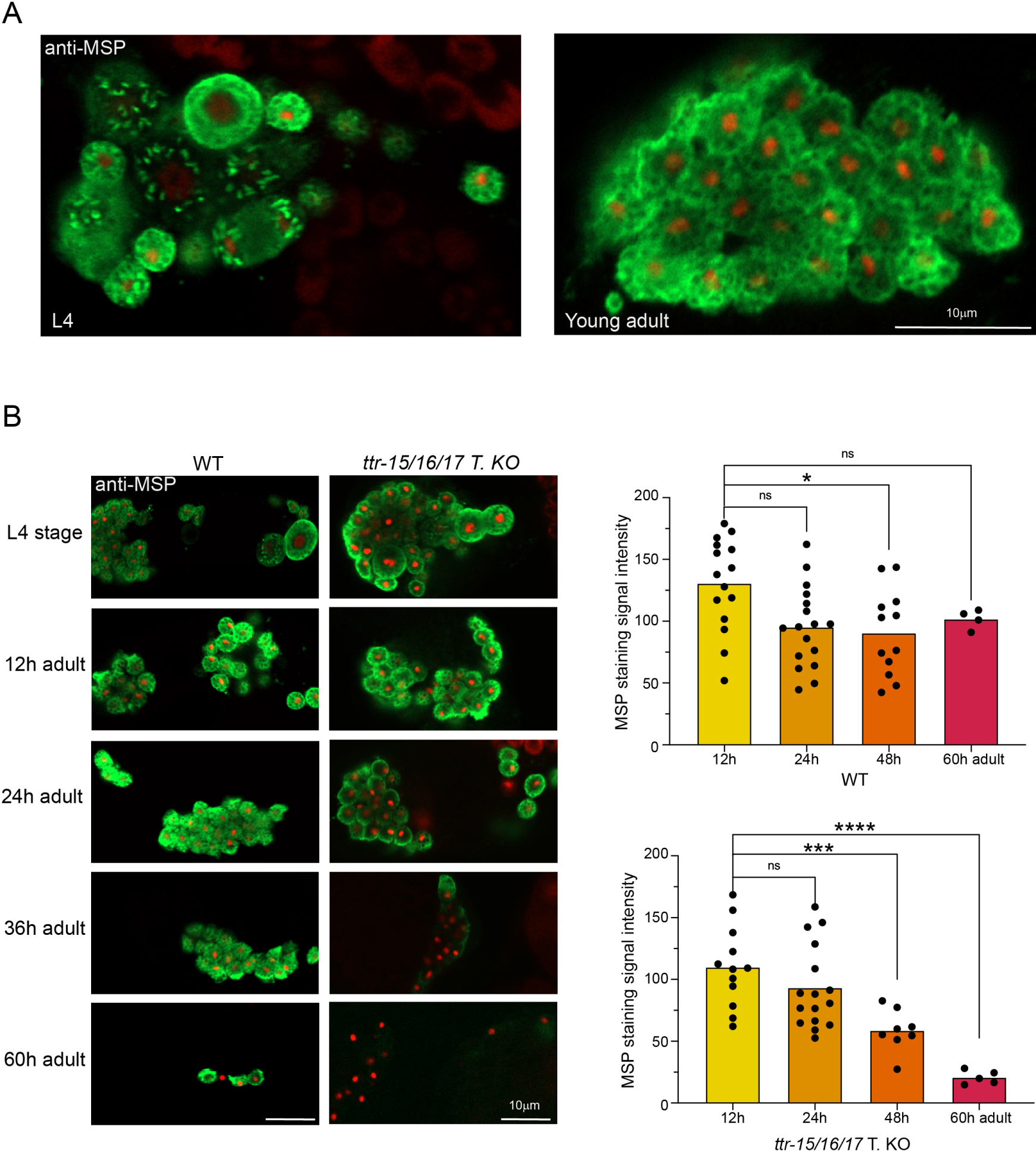
Accelerated Depletion of MSP from Sperm in Triple KO animals. A) Immunostaining of MSP on dissected gonads from wild-type animals. The MSP pattern exhibits a consistent distribution with previous reports. MSP is packed in fibrous organelles (FB) or diffused in the cytoplasm, depending on the stages of spermatogenesis in L4 animals (left). MSP is cytoplasmic in spermatozoa inside the spermatheca of young hermaphrodite adults (right). A vesicle-like structure with a hollow center is observed (arrow) and likely represents an MSP-containing double-membraned vesicle released from sperm. B) Immunostaining of MSP on dissected gonads from wild-type and triple KO animals. Genotype-matched, different-staged animals are processed in parallel. The MSP level drops rapidly from the second day in triple KO animals to a barely detectable level by the third day of adulthood. The intensity of MSP staining signals is quantified and shown on the right.

**Figure S4.**
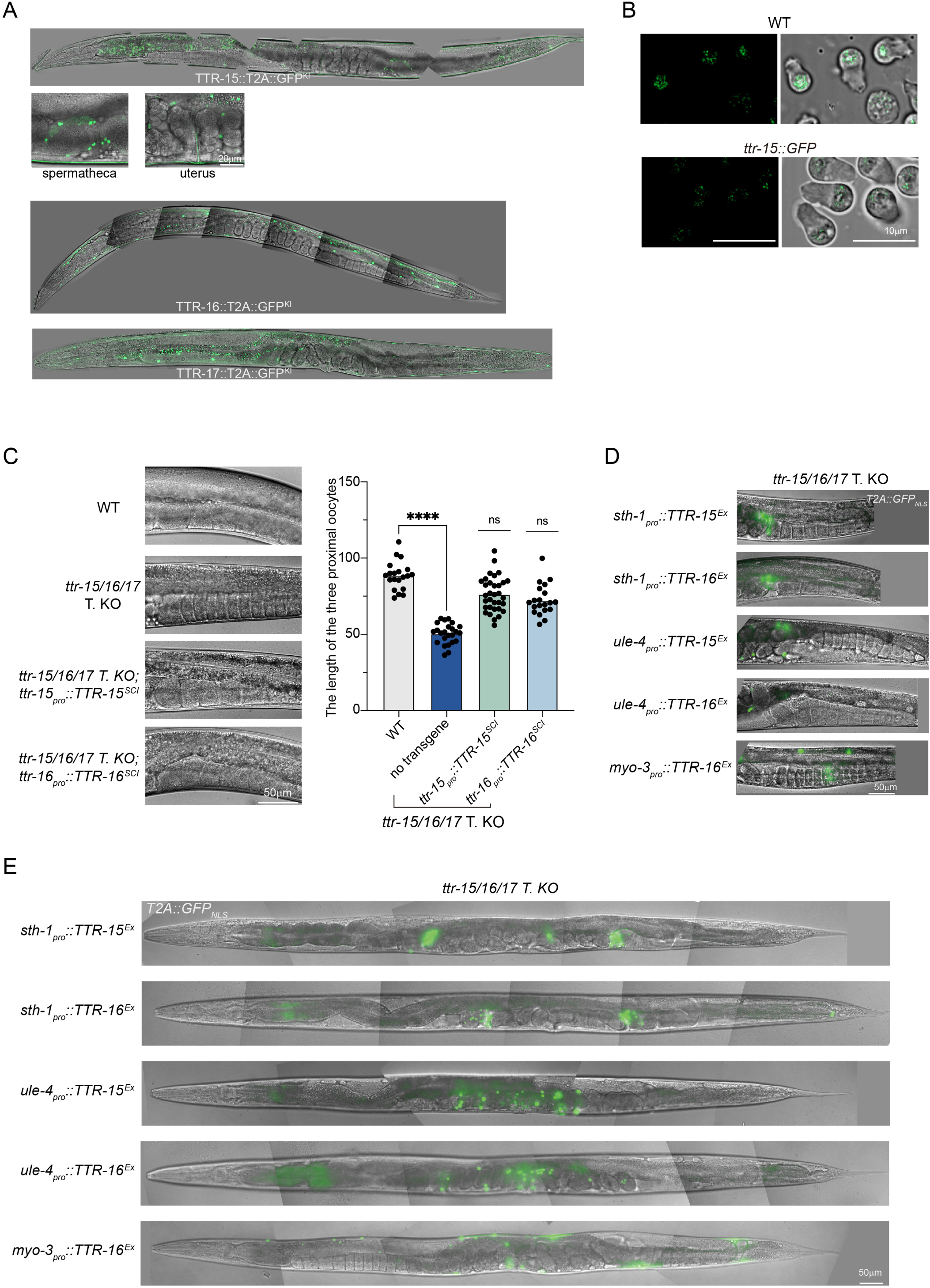
TTR-15, TTR-16, and TTR-17 are Critically Secreted from Somatic Gonads. A) Tissue expression of ttr-15, ttr-16, and ttr-17 labeled by endogenously tagged T2A::GFP-NLS. ttr-15 is primarily expressed in the spermatheca and uterus. ttr-16 is primarily detected in muscles and weakly in somatic cells of the gonads. ttr-17 is expressed in a similar pattern to ttr-16 but at much weaker intensity. B) Imaging of endogenously tagged TTR-15::GFP on dissected and washed male sperm; however, no GFP signals are detected. Controls are wild-type without the GFP tag. The green signals in the images are autofluorescence and captured by the high-power acquisition. C) The ovulation defects in triple KO animals are rescued by single copy insertion (SCI) transgenes of ttr-15 or ttr-16 minigene. Representative images focusing on gonads are shown (left). The length of the three most proximal oocytes is quantified (right). D) Representative images of tissue-specific expression of ttr-15 or ttr-16 transgenes in the triple KO background highlighting rescuing effects on blocked oocyte maturation. The expression of transgene is labeled by T2A::GFP-NLS.. E) Representative images of triple KO animals carrying tissue-specific expression of ttr-15 or ttr-16 transgenes highlighting the GFP localization. The presence of transgene expression is labeled by T2A::GFP-NLS. The ovulation in triple KO animals is well rescued by ttr-15 or ttr-16 expressed in the spermatheca.

